# Programming megakaryocytes to produce engineered platelets for delivering non-native proteins

**DOI:** 10.1101/2023.10.13.562311

**Authors:** Farhana Islam, Shwan B. Javdan, Mitchell R. Lewis, James D. Craig, Han Wu, Tara L. Deans

## Abstract

Platelets are anucleate cells naturally filled with secretory granules that store large amounts of protein to be released in response to certain physiological conditions. Cell engineering can endow platelets with the ability to deliver non-native proteins by modifying them as they develop during the cell fate process. This study presents a strategy to efficiently generate mouse platelets from pluripotent stem cells and demonstrates their potential as bioengineered protein delivery platforms. By modifying megakaryocytes, the progenitor cells of platelets, we successfully engineered platelets capable of packaging and delivering non-native proteins. These engineered platelets can offer flexible delivery platforms to release non-native proteins in a controlled manner upon activation when packaged into α-granules or deliver active enzymes to genetically alter recipient cells. Our findings highlight platelets as a promising tool for protein delivery in cell therapy applications.

## Introduction

The concept of using cells as a therapy to treat or prevent disease is becoming a reality with the engineering of bacteria, stem cells, and immune cells for this purpose. This approach is rapidly advancing the potential to transform medicine across disease areas ^1, 2, 3, 4, 5, 6, 7^. Cells have the natural ability to sense, integrate, and respond to dynamic changes in the body, making them attractive vehicles for targeting diseased cells to deliver various therapeutic agents ^8^. Synthetic biology offers state-of-the-art genetic tools to repurpose pathways for programming cells with customized performance that include switches, engineered receptors, and implementing Boolean logic for decision-making capabilities ^9, 10, 11, 12, 13, 14, 15, 16^. Using genetic tools and approaches in synthetic biology, cells can be enhanced with new functions to improve the delivery of non-native proteins to maximize therapeutic effects.

The type of cell used for developing cell therapies depends on the disease being treated and the desired function of the therapeutic cell ^17^. Platelets possess many unique characteristics that make them attractive candidates for the in vivo delivery of natural and synthetic payloads. They have an extensive circulation range in the body, accumulate at sites of injury, and naturally release biomolecules into the extracellular fluid upon activation ^18, 19, 20, 21^. In addition, platelets are anucleate cells, making them ideal therapeutic cells because they present no concerns about unwanted integration of foreign DNAs into the host genome. Therefore, platelets are well-equipped for mRNA and protein delivery.

Platelets are released into the bloodstream from a rare population of cells called megakaryocytes (MKs) that develop from hematopoietic stem cells (HSCs) ^22^. MKs can be successfully generated in vitro from pluripotent stem cells and HSCs, which enables the study of platelet development and applications of producing platelets in vitro ^23^. As MKs mature, the MK cytoskeleton is reorganized to promote long branched structures called proplatelets, that extend away from the MK cell body ^24^. Platelets can be released from the ends of proplatelets ^25, 26^, or produced from MK membrane budding ^27^. During MK maturation, proteins are packaged into α-granules that are transferred to platelets. Following platelet activation, α-granule contents are released into the surrounding environment to participate in a myriad of physiological processes ^28^.

In this study, we capitalize on the innate storage, trafficking, and release capabilities of platelets to engineer MKs during the cell fate process to express and package non-native proteins into platelets, creating engineered platelets. We show that these engineered platelets can be used for the targeted release of proteins upon platelet activation, and the genetic modification of recipient cells. Altogether, this work demonstrates a pipeline for creating a new cell therapy approach for delivering transgenic proteins. We envision that this body of work will facilitate studies leading to a better understanding of how proteins are packaged into platelets during the cell fate process, in addition to providing a new avenue for using platelets as a cell therapy.

## Results

### Production and scale-up of murine platelets in vitro

In developing a platform for engineering platelets as therapeutic cells, we considered their developmental pathway and the best approaches for genetic modification and expansion. Platelets are released from MKs, a rare cell population that accounts for less than 0.1% of nucleated cells in the bone marrow ^26, 29^. Due to this low number and their inability to divide ^30^, harvesting and genetically modifying MKs is impractical. While MKs are developed from HSCs, growing and expanding HSCs in vitro for creating stable genetically modified cells is challenging due to the small numbers initially harvested from patients, and the difficulty of maintaining HSCs beyond a few weeks in culture ^31, 32^. Pluripotent stem cells, on the other hand, can maintain their pluripotent state indefinitely in vitro or be induced to differentiate into any somatic cell type in the body ^33, 34, 35, 36^. An advantage to using pluripotent stem cells as a source for engineering and programming therapeutic cells is that they can easily undergo genetic modifications while maintaining a pluripotent state, and the genetically modified cells can be selectively expanded and differentiated into desired lineages ^37, 38^. Therefore, we chose mouse embryonic stem cells (mESCs) as our cell type to develop a pipeline for producing engineered platelets.

We previously showed that mESCs can be genetically programmed to express exogenous, transgenic proteins during differentiation to produce MKs expressing these proteins ^37^. Additionally, a study demonstrated that excessive bleeding in canines with hemophilia A could be improved with the delivery of platelets filled with human FVIII, a protein that plays a key role in clotting and hemostasis ^39^. This led to the hypothesis that platelets can be engineered and loaded with non-native proteins to be used as delivery vectors for modulating target cells by delivering their payloads. However, an important step for developing our cell engineering approach was to first create a pipeline to produce and scale up the production of platelets in vitro from mESCs (Fig. 1a). A common method for differentiating mESCs into HSCs involves plating them on a layer of OP9 cells for 5 days ^40^. After mESCs commit to the hematopoietic lineage, thrombopoietin (TPO) is added to the culture. TPO signaling in the bone marrow is essential for HSC survival, proliferation, and differentiation into MKs, making TPO a commonly used cytokine for stimulating and differentiating HSCs to become MKs ^26, 41, 42, 43^. TPO stimulation results in the expression of CD41, a key surface marker indicating MK lineage commitment. In our pipeline, after culturing with TPO for 7 days, cells were stained with CD41 antibody and imaged (Fig. 1b). When culturing mature MKs in vitro, platelets can be shed from MKs and harvested from the media. CD41 stained MKs (Fig. 1c and Supplemental Fig. 1a) and platelets (Fig. 1d and Supplemental Fig. 1b) were collected and run on a flow cytometer confirming that CD41 expression increased after differentiation when compared to mESCs. A characteristic of achieving functional platelets is their ability to activate when exposed to thrombin and ADP, which initiates platelets to express a large amount of P-selectin ^44^. Studies have shown that adding ADP to the reaction plays an important role in thrombin-induced platelet aggregation and activation ^45, 46^. After harvesting platelets from MKs produced in vitro, they were exposed to thrombin and ADP. P-selection expression was monitored over time and the maximum activation was observed after 3 hours (Fig. 1e), where a little over 80% of the platelets were activated (Supplementary Fig. 2). It is notable that the activation of in vitro produced platelets takes longer than what is observed in vivo, likely because the in vitro conditions lack dynamic flow and other components that naturally trigger platelet activation following wounding in vivo ^47, 48, 49^.

**Fig. 1.**
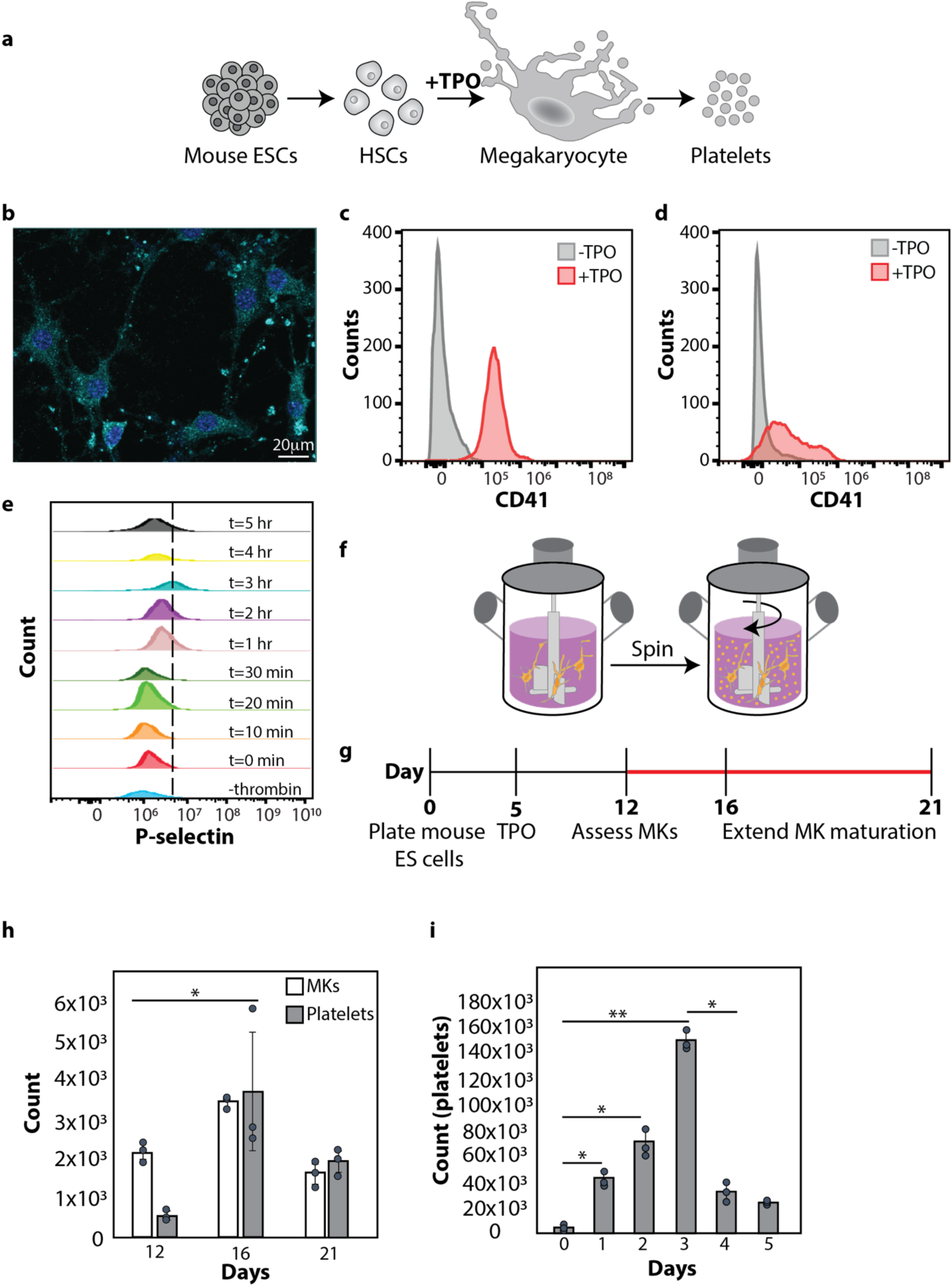
Production of murine MKs and platelets from mESCs in vitro. **a** Schematic representation of differentiating mESCs to produce platelets in vitro. **b** Fluorescent image after 12 days of differentiation stained with MK lineage marker, CD41 (teal) and nuclei stained with DAPI (blue). Scale bar is 20μm. **c** Flow cytometry of undifferentiated mESCs (-TPO, grey) and differentiated MKs (+TPO, red) stained with CD41. **d** Flow cytometry of undifferentiated mESCs (-TPO, grey) and differentiated platelets (+TPO, red) stained with CD41. **e** Timeline of mouse platelet activation after exposure to thrombin/ADP. Dotted line indicates the boundary of platelets not activated (left of line) and activated (right of line). **f** Spinner flask bioreactors to scale up the production of platelets in vitro. **g**. Scaling up mouse platelet production in vitro by extending the MK maturation beyond 12 days (red line). **h** MK maturation optimization. Flow cytometry comparing the number of MKs and platelets produced by extending the MK maturation beyond 12 days. The number of cells represents the mean from three independent experiments. The error bars represent the standard deviation of the mean. *p ≤ 0.05. **i** Time in spinner flask bioreactor. Flow cytometry of the number of platelets produced over time in the spinner flask bioreactors. The number of platelets represents the mean from three independent experiments. The error bars represent the standard deviation of the mean. *p ≤ 0.05, **p ≤ 0.01.

As with all cell therapies, large numbers of cells are required for efficacy. Conventional 2D culturing systems are insufficient for producing the high number of platelets required for transfusion. For human transfusions, a typical therapeutic dose is at least 3x10^11^ platelets for an adult patient, and 3x10^6^ platelets per mouse in a mouse study ^50, 51^. In vivo, intravascular shear forces generated by blood flow play a crucial role in the release of platelets from MKs ^52, 53^. To emphasize the scalability of our pipeline, production was scaled up using a spinner flask bioreactor (Fig. 1f). To explore the ideal number of MKs to produce the highest number of platelets, we first assayed MK numbers using our original 12-day differentiation protocol and then extended the protocol up to 21 days to determine if this extra culturing time increased the number of MKs and whether more mature MKs could produce a higher number of platelets (Fig. 1g). We found that extending the differentiation protocol to 16 days increased the number of MKs and platelets from the 2D culture (Fig. 1h). Therefore, after 16 days of differentiation and maturation, cells were transferred to a spinner flask bioreactor, and the number of platelets was assessed over a 5-day period. We found that maturing MKs for 16 days and transferring them to a spinner flask bioreactor for 3 days yielded the greatest number of platelets (Fig. 1i). This scaling up approach is not restricted to mESCs and can also be effectively applied to human cell lines. We demonstrated this by evaluating MEG-01s, a human megakaryoblast leukemia cell line. MEG-01s are an immature MK cell line that can be matured with exposure to TPO to resemble natural MKs (Supplementary Fig. 4a). When MEG-01s mature, they produce membranous extensions, increase DNA content, and release platelet-like particles (PLPs) that have characteristics resembling human platelets (Supplementary Fig. 4b) ^54, 55, 56^. Similar to mESCs, human MEG-01 cells could be matured and put into a spinner flask bioreactor to produce larger numbers of PLPs compared to harvesting them from 2D cultures (Supplementary Figs. 3 and 4). Together these studies show that using progenitor stem cells as a cell source for producing platelets in vitro is promising and has the potential to be scaled up using spinner flask bioreactors.

### Loading non-native proteins into platelets

By generating platelets from mESCs in vitro, we created the opportunity to enhance platelets with new functions by engineering protein cargos. As proof of concept, on day 10 of the mESC differentiation pipeline, MK progenitor cells were transfected with a plasmid that constitutively expresses GFP, which was used as a proxy to establish proof of concept for other protein cargos since it is not toxic and easily detected (Fig. 2a and Supplementary Fig. 5a). GFP expression was observed throughout the cytoplasm 48 hours post-transfection (Fig. 2b). The native protein, von Willebrand factor (VWF), was also observed further indicating that our engineered MKs were mature (Fig. 2b) ^57^. GFP expression was quantified using flow cytometry (Supplementary Fig. 5) that showed the transfection efficiency to be around 19% (Supplementary Fig 5e). For successfully transfected cells, the bell-shaped GFP expression curve represents that most cells have moderate level of GFP loading. A smaller proportion of MKs express very high or low levels of GFP compared to non-transfected MKs (Fig. 2c). Next, efficient loading of MK derived platelets with GFP was tested by flow cytometry and compared to non-transfected cells, confirming that platelets expressed GFP (Fig. 2d). This was also seen in the human cell line, MEG-01s, where GFP could be expressed after being transfected with a plasmid constitutively expressing GFP (Fig. 2e). Similar to mESCs, MEG-01s expressing GFP were matured into MKs in the presence of TPO and GFP expression was observed throughout the cytoplasm 48 hours after transfection (Fig. 2f). PLPs were harvested from transfected MEG-01 derived MKs and GFP expression was also observed in them (Fig. 2g). Flow cytometry confirms the maturation of MEG-01s to MKs in the presence of TPO with the increased expression of the MK lineage marker, CD41 (Fig. 2h). GFP expression was also compared to non-transfected cells using flow cytometry and there is a clear shift confirming that MKs express GFP (Fig. 2i).

**Fig. 2.**
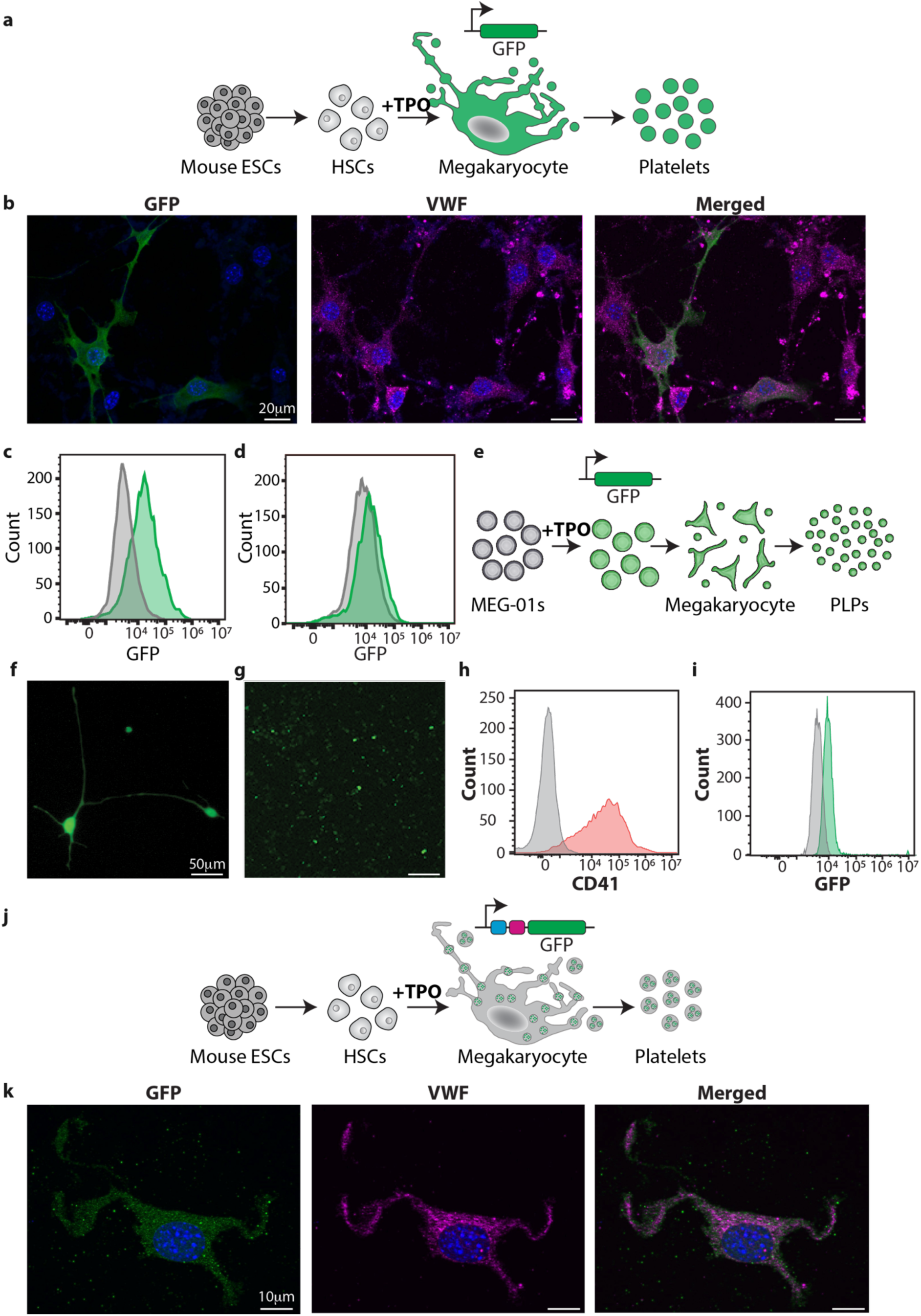
Programming mouse MKs to load non-native protein in platelets. **a** Schematic representation of programming mouse MKs during differentiation to load non-native proteins in platelets in vitro. **b** Fluorescent image after 12 days of differentiation expressing GFP (green) in MKs, Von Willebrand factor (VWF, pink) and nuclei stained with DAPI (blue). All scale bars at 20μm. **c** Flow cytometry of untransfected (grey) and transfected (green) MKs with GFP. **d** Flow cytometry of platelets from untransfected (grey) and transfected (green) MKs with GFP. **e** Schematic representation of programming MEG-01 cells to express GFP to be packaged into platelets during MK maturation. **f** Fluorescent image of MEG-01s transfected with GFP and matured to MKs. Scale bar is 50μm. **g** Platelets obtained from MKs derived from MEG-01s expressing GFP. Scale bar is 50μm. **h** Flow cytometry of undifferentiated MEG-01s (-TPO, grey) and differentiated (+TPO, orange) MKs stained with the MK lineage marker, CD41. **i** Flow cytometry of MKs not transfected with GFP (grey) and MKs transfected with GFP (green). **j** Schematic representation of GFP fused to peptides for targeting GFP to α-granules. **k** Fluorescent images of GFP targeted to the α-granules (green), Von Willebrand factor (VWF, pink) and nuclei stained with DAPI (blue). All scale bars are 10μm.

Platelets secrete proteins by loading them into α-granules that are released following activation ^28^. To engineer platelets with controlled release capabilities of our transgenic proteins, we first wanted to demonstrate that they could be packaged into α-granules in MKs in our pipeline of making platelets from mESCs in vitro. Similar to previous work, we packaged GFP into MK α-granules by fusing GFP to a short peptide sorting signal derived from the human cytokine RANTES to its 5’ end ^58^ (Fig. 2j and Supplementary Fig. 6). Without the peptide fused to GFP, the protein is evenly distribution throughout the MK cytoplasm (Fig. 2b). Conversely, fusing the peptide to GFP targets it to the α-granules and displays a different distribution of GFP in the cytoplasm of mouse MK cells, with a dotted appearance throughout the cytoplasm (Fig. 2k). VWF is naturally synthesized in mature MKs and is primarily stored in α-granules that are trafficked down MK proplatelets and transferred to platelets ^59^. Staining for VWF and GFP confirms their co-localization within the MK α-granules (Fig. 2k). When observing the GFP distribution in the human MEG-01 cell line, GFP expression was also noted to be unevenly distributed in the cytoplasm when targeted to the α-granules (Supplementary Fig. 7).

### Engineered platelets for the controlled release by targeting non-native proteins to α-granules

Because proteins are released from α-granules upon platelet activation, we reasoned that the packaging can be modified to include non-native proteins for therapeutic protein delivery. To evaluate this mode of delivery, we assessed the secretion of secreted alkaline phosphatase (SEAP) targeted and non-targeted to α-granules over a 9-hour period of time. We chose to evaluate SEAP in this context because it is a naturally secreted protein, and is widely used as a secreted protein reporter ^60^. Furthermore, since SEAP is a naturally secreted protein, we wanted to see whether we could keep it in the α-granules over time. To load mouse platelets with SEAP, MKs were transfected on day 10 of our differentiation pipeline, and on day 12, platelets loaded with SEAP were harvested and resuspended in fresh culture media to study its release under thrombin/ADP-induced activation using a calorimetric SEAP quantification assay. As expected, platelets containing SEAP not targeted to the α-granules (Fig. 3a, Supplementary Fig. 8a) readily released it into the media (Fig. 3c). However, when the RANTES α-granule peptide was added to the SEAP coding sequence (Fig. 3b, Supplementary Fig. 8b), SEAP was maintained in the cell over time and was only released when the platelets were activated by thrombin/ADP (Fig. 3c). We observed a sharp decrease in the α-granule targeted SEAP 3 hours after activation. We believe that this is attributed to the addition of thrombin and ADP to the culture media, which causes an immediate change of phenol red from pink to colorless, indicating a shift to an acidic pH. SEAP released from the platelets is exposed to this acidic environment for an extended period (3-4 hours), which likely led to its degradation. Consistent with the human MEG-01 cell line in our studies, we see similar results of consistent secretion of SEAP without being targeted to the α-granules and when SEAP is targeted to the α-granules, we only see SEAP released into the media after the PLP activation with thrombin/ADP (Supplementary Fig. 10).

**Fig. 3.**
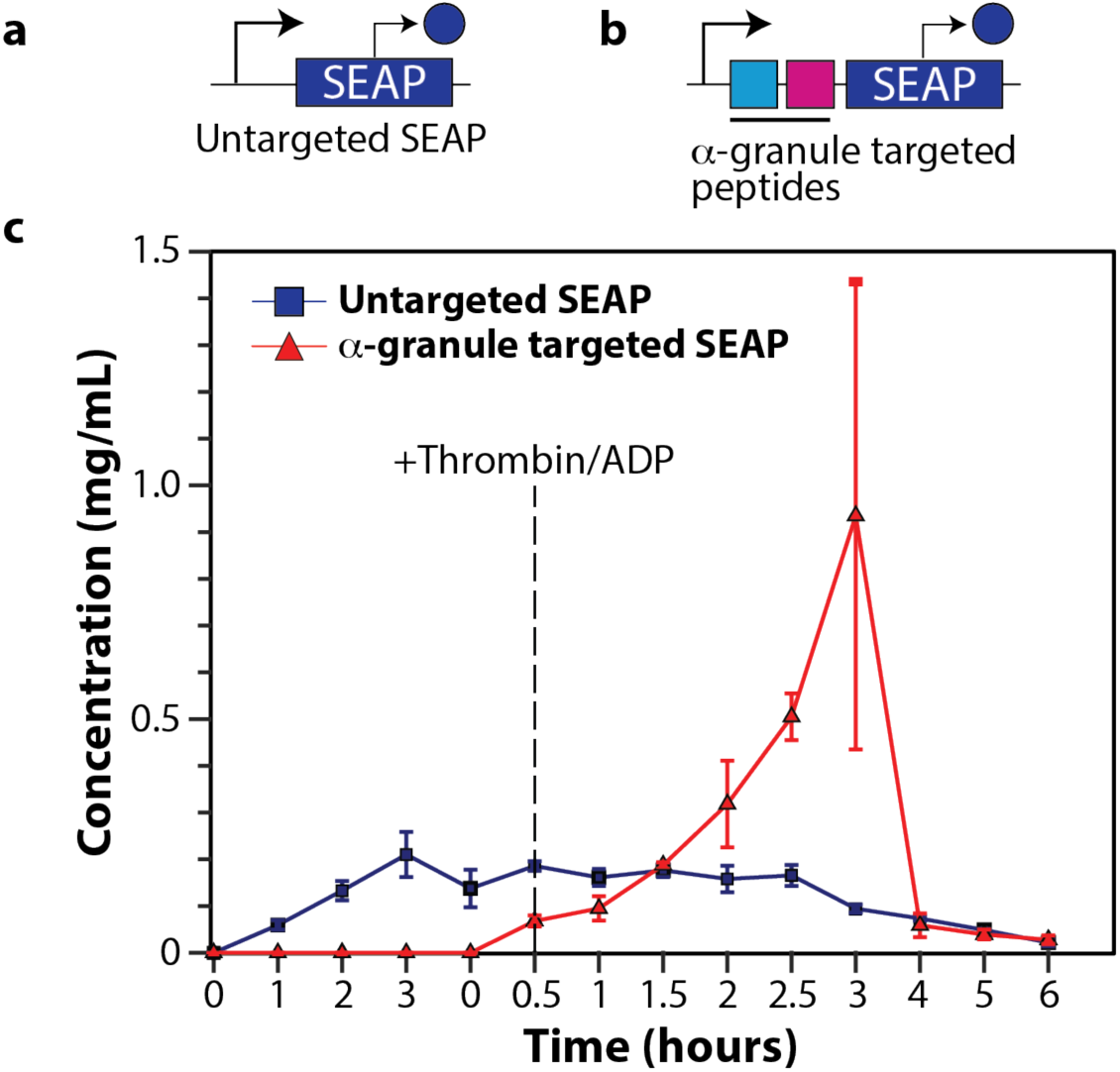
Targeting SEAP proteins to α-granules for their activated release in mESCs. **a** Schematic representation of constitutively expressing SEAP not targeted to the α-granules. **b** Schematic representation of SEAP fused to peptides for targeting to α-granules. **c** Non-α-granule-targeted SEAP (blue) over time. SEAP targeted to the α-granules (red) and its release upon activation with the addition of thrombin and ADP (dotted line). Experiments were repeated independently at least three times with similar results. Error bars represent the standard deviation of the mean.

### Engineered platelets can deliver functional proteins into recipient cells

To investigate a second mode of delivery that did not involve pre-packaging of non-native proteins into α-granules, the delivery of bioactive proteins loaded into the cytoplasm of engineered platelets was investigated. Cre recombinase is an enzyme that catalyzes the site-specific recombination of DNA between loxP sites ^61^. We chose to load platelets with Cre recombinase because we wanted to test whether the transgenic protein in engineered platelets would be functional after being delivered to recipient cells. If Cre recombinase is functional, it will perform a genetic alteration and remove specific DNA sequences between loxP sites. To evaluate platelet-dependent delivery of Cre recombinase, we used an HEK293 reporter cell line that contains loxP-GFP-RFP. In this cell line, the GFP coding sequence is flanked by loxP sites and is positioned upstream of RFP (Fig 4a). In the absence of Cre recombinase, the cells express GFP, and translational stops, preventing the expression of RFP. When Cre recombinase enters these cells, the enzyme excises the DNA fragment between the loxP sites, thereby allowing the cells to express RFP instead of GFP. However, Cre recombinase is normally enriched in the nucleus and therefore not readily available in the cytoplasm of MKs to be packaged into platelets. Therefore, the Cre fused with the estrogen receptor (ER) was used because it maintains the modified Cre-ER in the MK cytoplasm. Cre-ER can be activated by 4-hydroxy tamoxifen (4-OHT) to transfer it to the nucleus of the reporter cells. Upon activation, Cre-ER translocates to the nucleus ^62^, enabling Cre to perform the DNA excision of GFP and the stop sequence. We reasoned that cytoplasmic Cre-ER would be readily packaged into platelets if it was expressed in MKs.

**Fig. 4.**
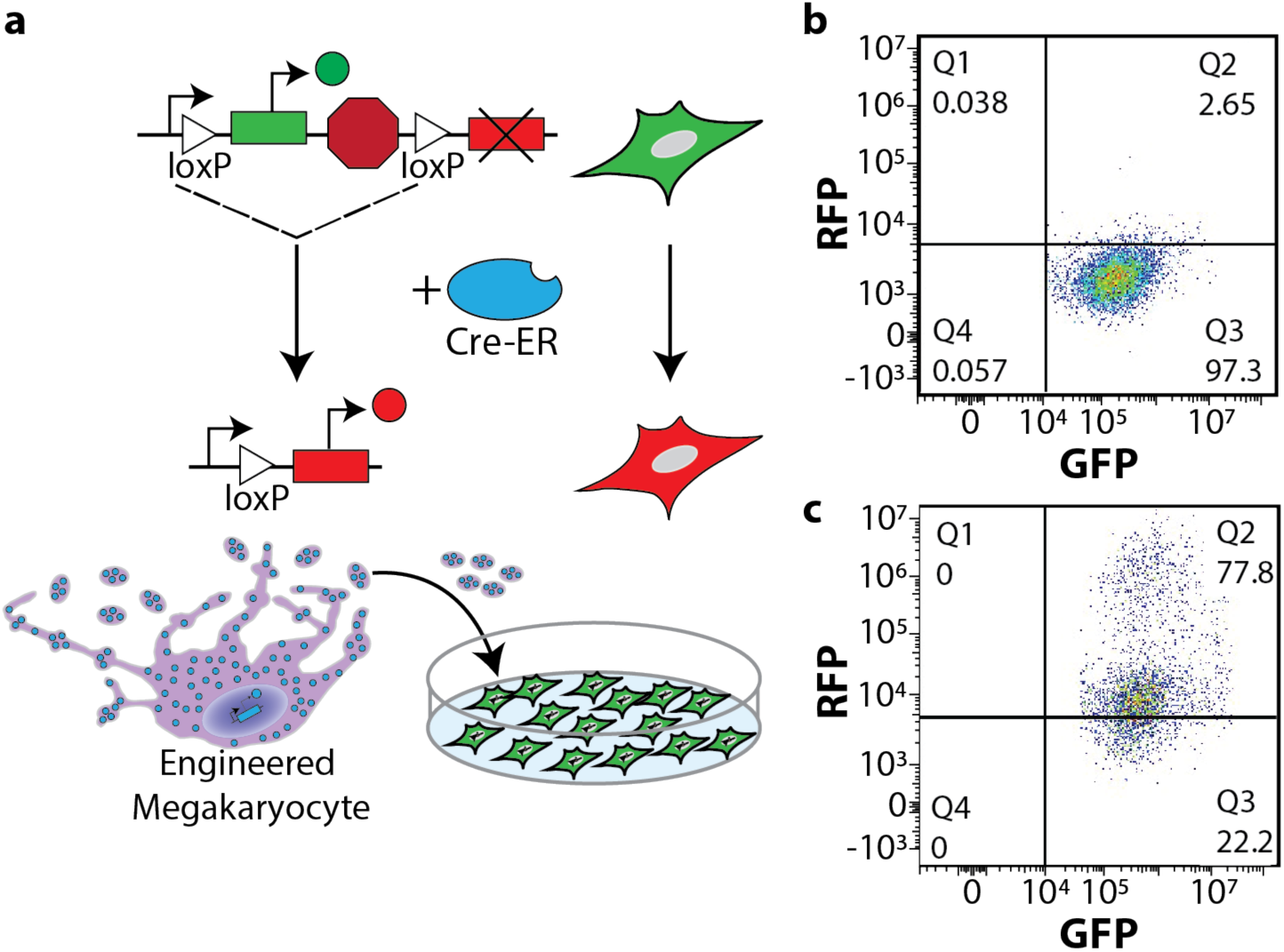
Loading mouse platelets with Cre recombinase genetically alters recipient cells after delivery. **a** Schematic representation of LoxP-GFP-RFP Cre reporter HEK293 cell line. In the presence of Cre recombinase (blue) the HEK cells turn from green to red with the removal of GFP and the stop sequence. Platelets loaded with Cre were harvested and co-cultured with the HEK293 reporter line. **b** Flow cytometry of the reporter HEK293 cell line before Cre-loaded platelets were added to the culture. **c** Flow cytometry of the reporter HEK293 cell line after Cre-loaded platelets were added to the culture.

Because our transient transfection efficiency for plasmids expressing GFP in MKs was approximately 19% (Supplementary Fig. 5e), we needed a way to monitor the transfection efficiency of the Cre-ER plasmid to ensure successful transfections before harvesting platelets. To accomplish this, we built a bicistronic message containing the GFP and Cre-ER coding sequences separated by a P2A peptide sequence (Supplementary Fig. 11). This additional GFP to the non-transfected HEK293 reporter cells should not impact the evaluation of Cre recombinase function because such recombination is determined by the RFP expression in the HEK293 reporter cells. To load mouse platelets with Cre-ER, MKs were transfected on day 10 of our differentiation pipeline and successful transfections were confirmed by the expression of GFP 48 hours later. At this time platelets were harvested from the media, washed, and co-cultured with the HEK293 reporter line. Subsequently, 4-OHT was added to the co-culture to facilitate Cre-ER translocation into the HEK293 nuclei and activate RFP expression. Flow cytometry revealed that the HEK293 reporter cells mainly expressed GFP before co-culturing with Cre-ER loaded platelets (Fig. 4b and Supplementary Fig. 13d) and a drastic change in RFP expression 48 hours after co-culture and 4-OHT addition (Fig. 4c and Supplementary Fig 13e) compared to HEK293 reporter cells alone (Fig. 4b). Since the collection of the Cre-ER filled platelets involved removing platelets from the culture media of differentiating cells and washing them, we are confident that the Cre-ER presented to the HEK293 reporter cells is from the platelets and not from Cre-ER that may have been present in the media from dying MKs. We believe that the engineered platelets were engulfed by the HEK293 reporter cells after settling on them during the 48-hour co-culturing period, or that they released small particles containing Cre-ER that were engulfed by the HEK293 cells ^63^. These results were also seen in the human MEG-01 cell line where PLPs filled with Cre-ER delivered to HEK293 reporter cells flipped the expression from GFP to RFP after 48 hours of incubation (Supplementary Fig. 14).

## Discussion

Platelets are anucleate blood cells that circulate throughout the body with diverse roles in hemostasis, wound healing, angiogenesis, inflammation, and clot formation. They are naturally filled with secretory granules that store large amounts of bioactive proteins and are released upon activation. These unique characteristics make them attractive candidates to repurpose their protein packaging during hematopoiesis to create new delivery vehicles for cell therapies. This study genetically modified megakaryocytes, the progenitor cells of platelets, during the cell fate process to load platelets with non-native proteins. We show these engineered platelets can be loaded with various kinds of proteins, and these proteins can be released by different delivery mechanisms.

Scaling up the production of autologous cells as a starting material for therapies remains challenging due to the high variability of patient cells that must successfully undergo genetic alterations and retain the ability to respond to signals in vivo when reinfused ^64^. Using pluripotent stem cells as an initial cell source is advantageous because these cells can be expanded indefinitely, and they have the potential to become any somatic cell in the body. We and others have shown that murine and human platelets can be successfully made in vitro from pluripotent stem cells ^37, 52, 55, 65, 66, 67^. Our results demonstrate that murine pluripotent stem cells can be differentiated down the hematopoietic lineage to generate large numbers of platelets in vitro by scalable methods. As a proof of concept for scaling up platelet production in our studies, we used spinner flask bioreactors that showed significantly improved platelet counts compared to harvesting platelets without turbulent flow, which is similar to other studies ^52^. Specifically, we found that MKs taken from a single well of a 12-well plate can improve platelet production over 50-fold compared to without turbulent flow (Fig. 1). Establishing this pipeline of platelet production in vitro offers an approach to scale up pluripotent stem cells as a source for establishing cell therapies. Furthermore, there is a significant advantage to using pluripotent stem cells (e.g. mESCs and human induced pluripotent stem cells (iPSCs)) as a starting material because mESCs can be derived from the same genetic background as disease models, and human iPSCs can be derived from individual patients. Both of which are not prone to genetic instabilities associated with the long-term culture of immortalized cell lines. Furthermore, efforts are currently being made to genetically edit human leukocyte antigen (HLA) in human iPSCs to create a universal cell line that would be compatible with a wide range of patients ^68^. Altogether, pluripotent stem cells are an ideal initial cell source for establishing a pipeline for generating cell therapies.

In addition to pluripotent stem cells having the capacity to grow indefinitely, they can easily undergo genetic modifications while maintaining a pluripotent state and be selectively expanded. Furthermore, cells can be genetically modified during the differentiation process to obtain desired lineages with those modifications. In this study, we showed that MKs can be genetically modified during the cell fate process to express proteins to be packaged into platelets (Fig. 2). While the efficiency of this approach was enough for the proof of concept at smaller scales, this process could be improved by making stable pluripotent stem cell lines with the desired genetic modifications and differentiating those cells to become MKs. However, one challenge that may arise with this approach is that the expression of certain proteins that may be desired for therapeutic delivery could negatively impact the cell fate process and prevent the robust production of modified MKs and engineered platelets. One possible solution to this challenge is to use synthetic gene circuits that can tightly control gene expression to keep genes in the off state until turned on by an inducer molecule once the cells reach the MK stage ^69^. A second possible solution is the use of MK specific promoters that will turn on the expression of the desired gene only when the cells have reached the MK stage of development. An additional challenge to programming pluripotent stem cells is transgene silencing ^70^. Significant efforts are underway to better understand this phenomenon for developing approaches to mitigate it ^71, 72, 73^.

By genetically modifying murine and human MKs during the cell fate process, we demonstrated two distinct methods of loading and delivering non-native proteins using platelets. One by packaging the non-native protein into α-granules and their release upon platelet activation, and the other by keeping the non-native protein in the MK cytoplasm to load into platelets, which can be engulfed by the target cells. Consistent with other reports ^58^, our data shows that non-native proteins can be packaged into α-granules in MKs by fusing the protein to the sorting signal of RANTES. To demonstrate the targeted release from platelets using this delivery method, we fused SEAP to the sorting signal of RANTES and show that SEAP is only released into the media after platelet activation with exposure to thrombin. Recently a study highlighted protein localization in MKs and their transport during the formation of platelets, demonstrating that proteins can be trafficked to platelets by different mechanisms ^74^. The second method of delivery showed the loading of a nuclear protein from the MK platelets by fusing it with estrogen receptor. Previous work has shown that platelets can be engulfed by immune cells ^75, 76, 77^ and their microparticles can deliver nucleotides and proteins to other cells ^63, 78, 79^, suggesting that this type of delivery can be used for modifying neighboring cells. In this study, we show that Cre recombinase remains active after being packaged into platelets without granule targeting, and when delivered to recipient cells harboring loxP sites within their genome, perform homologous recombination to remove the DNA sequence between the loxP sites (Fig. 4). Cre is likely transferred from platelets to the HEK293 cells either through platelet engulfment or microparticle delivery.

Our proof-of-concept studies reveal an innovative and flexible approach to deliver therapeutic protein payloads to recipient cells using engineered platelets. We show that the delivery of these proteins is diverse and can be used to genetically program target cells, to continuously release proteins over time, or for the therapeutic protein to remain localized in α-granules to be released upon platelet activation. These results highlight the utility of programming MKs with therapeutic proteins to be packaged into platelets that can be engineered with a variety of release profiles depending on the therapeutic need and offer an exciting entry point to develop new therapies for delivering therapeutic proteins and programming cells in vivo. We imagine that engineered platelets can be used for many therapeutic applications including the delivery of metabolic enzymes that may be missing in a patient, delivering payloads to tumors to stunt their growth, modifying immune cells to enhance or dampen their response, or prevent cardiovascular disease by mitigating the buildup of plaques to prevent atherosclerosis. To enhance many of these applications, these delivery methods could be improved by engineering peptides, nanobodies, or chimeric receptors on the surface of platelets to improve targeting of specific cell types and tissues for delivery.

## Methods

### Cell lines and differentiation

Detailed protocols for growing and differentiating mESCs and MEG-01s have been previously published by our lab ^37, 55^. In short, D3 mouse embryonic stem cells (mESCs) were obtained from ATCC (#CRL-1934). Neomycin resistant mouse embryonic fibroblasts (MEFs) were used as a feeder layer for the ESCs (Fisher Scientific #PMEF-NL-P1) and Mitomycin C (Fisher Scientific #BP2531-2) treated before co-culturing with ESCs. The mESCs were maintained in high glucose knockout DMEM (ThermoFisher Scientific #10829018) supplemented with 15% final concentration ES certified FBS (ThermoFisher Scientific #10439024), 1% final concentration nonessential amino acids (ThermoFisher Scientific #11140050), 1 mM L-glutamine (ThermoFisher Scientific #25030-081), 0.1 mM 2-mercaptoethanol (ThermoFisher Scientific #21985023), 10 ng/mL LIF (ThermoFisher Scientific #PMC9484) and 1% final concentration penicillin/streptomycin (ThermoFisher Scientific #15140-122). The MEFs were expanded on 0.1% final concentration gelatin coated plates and grown in high glucose DMEM medium (ThermoFisher #11965-092) supplemented with 10% final concentration FBS (ThermoFisher Scientific #A5256701), 1% final concentration nonessential amino acids, 1mM L-glutamine, and 1% final concentration penicillin/streptomycin. For the differentiation studies of mESCs, OP9 cells (ATCC #CRL-2749) were seeded in one well of a 12-well plate and grown in MEM α, no nucleoside medium (ThermoFisher Scientific #12561-056) containing 20% final concentration FBS (ThermoFisher Scientific #A5256701), and 1% final concentration of penicillin/streptomycin. Recombinant mouse thrombopoietin (TPO) (VWR #315-14-50UG) was added at 20 ng/mL until day 8 of differentiation where it was then decreased to 10 ng/mL until cells were harvested for analysis ^37^. The MEG-01 cells (ATCC #CRL-2021) were grown in RPMI 1640 medium (ThermoFisher Scientific #11875093) that contains 10% final concentration FBS and 1% final concentration penicillin/streptomycin. MEG-01 cells were matured in growth medium containing 100 ng/mL recombinant human TPO (VWR #10773-60) for 48 hours. The HEK293 reporter cell line (FisherScientific #NC1639588) was grown in high glucose DMEM medium (ThermoFisher #11965-092) supplemented with 10% final concentration FBS, and 1% final concentration penicillin/streptomycin. All cell lines were grown in a humidified 5% CO_2_, 37 °C incubator.

### Mouse platelet, MEG-01 platelet like particle (PLP), and MK collection

Once the cells have differentiated and matured in the presence of TPO (day 12 of mESC differentiation and day 3 of MEG-01 differentiation), platelets and PLPs were harvested by transferring the medium from the respective cultures to separate 15 mL conical tubes. Cells were centrifuged at 100xg for 5 minutes to remove larger cells and debris, followed by centrifugation at 1200xg for 15 minutes to pellet and isolate the platelets and PLPs.

Mouse adherent MKs were detached from the culture dish using 0.25% trypsin (ThermoFisher Scientific #25200056) and neutralized with high glucose DMEM medium supplemented with 10% final concentration FBS, and 1% final concentration penicillin/streptomycin. MEG-01 MKs were scraped with a cell scraper to remove cells. Once cells were removed from the tissue culture dish, cells were transferred to a separate 15 mL conical tube. Cells were centrifuged at 100 xg for 5 minutes to pellet and isolate the MKs.

### Platelet activation study

On day 12 of mESC differentiation, platelets were collected and resuspended in fresh OP9 medium with 40 μl thrombin and 3.42 μl of ADP (Millipore Sigma #A-2754) from the stocks were added to 1 ml OP9 media. This means the concentration of ADP in 1 ml of medium was 191.81 uM and thrombin was 1.92 U/ml that we used for this study. 2 U/mL of mouse thrombin (Fisher Scientific #50-203-6453) was added to the resuspended cells. Following activation, platelets were collected at various time points and washed with PBS by centrifugation at 1200 xg. Platelets were resuspended in 100 μL of PBS containing 3% BSA for antibody labeling and flow cytometry analysis.

### High throughput production of platelets and PLPs

A streamlined, two-phase approach was developed to scale up platelet production. First, producing mature MKs in static culture (2D tissue culture plate), followed by applying shear stress in a spinner flask bioreactor to stimulate platelet production.

Scaling up mouse platelets started with optimizing the process by first assessing whether maturing MKs beyond 12 days during the differentiation method would improve the number of platelets produced using a spinner flask bioreactor. To test this, mESC-derived MKs were grown beyond 12 days. Cells were passed on days 12 and 16 onto the fresh OP9 layer with media supplemented with 10 ng/mL TPO. Since we found that extending the maturation of MKs to 16 days improved platelet yields, we moved these cultures into a 125 mL glass spinner flask bioreactor (VWR #22877-073) containing 99 mL of fresh OP9 culture medium. Specifically, on day 16 MKs and platelets were harvested using 0.25% trypsin, deactivated with complete DMEM and washed with PBS. The cell pellet was resuspended in 1 mL fresh OP9 medium and transferred to s spinner flask bioreactor containing 90 mL of OP9 media. Shear stress was applied using a dura-mag magnetic stirrer (VWR #30624-012) set to 100 rpm ^80^. The spinner flask bioreactor was incubated at 37°C for 7 days with 500 μL samples collected every 24 hours. After each sampling, an equal volume of OP9 growth medium was added to maintain a consistent volume throughout the study. MKs and platelets from the samples were isolated using the previously described methods and analyzed by flow cytometry.

To scale up PLPs, MEG-01 cells were matured in 100 ng/mL TPO for 72 hours in static culture, then the cells were scraped using a cell scraper to dislodge the MKs. The entire population of cells in the well were transferred to a 125 mL glass spinner flask bioreactor (VWR #22877-073) containing 99 mL of fresh MEG-01 culture medium. Shear stress was applied using a dura-mag magnetic stirrer (VWR #30624-012) set to 100 rpm. The spinner flask bioreactor was incubated at 37°C for 7 days with 500 μL samples collected every 24 hours. After each sampling, an equal volume of MEG-01 growth medium was added to maintain a consistent volume throughout the study. MKs and PLPs from the samples were isolated using the previously described methods and analyzed by flow cytometry.

### Design of plasmids

TLD_010 (Addgene #19696) was used to constitutively express GFP in MKs. Due to complexity, the plasmid that expresses Cre-ER was constructed in parts by gBlock synthesis of DNA fragments (Integrated DNA Technologies) and assembled using standard molecular biology techniques including restriction enzyme digest (NEB) to cut the DNA fragments, PureLink™ Quick Extraction Kit (ThermoFisher K210025) to extract DNA from 1% agarose (Fisher Scientific BP1356-100) gels and T4 ligase (Thermo Fisher Scientific 15224017) to ligate the pieces together. Information on the cloning and deposition of plasmids constructed during this study can be found in Supplementary Fig. 10 and Supplementary Table 2. CMV-SEAP (Addgene #24595) was used to constitutively express SEAP in MKs. The α-granule targeted EGFP plasmid was made by gBlock synthesis of the short peptide sorting signals derived from the human cytokine RANTES and cloned into the TLD_010 plasmid to target the GFP to α-granules (Supplementary Fig. 6). The α-granule targeted SEAP plasmid was made by gBlock synthesis of the short peptide sorting signals derived from the human cytokine RANTES and cloned into the CMV-SEAP plasmid (Supplementary Fig. 6). Information on the cloning can be found in Supplementary Table 2. To improve translation efficiency for expressing SEAP in the mESCs, the Kozak sequence 5’-GCCACCATG-3’ was purchased as oligonucleotides from the University of Utah DNA/Peptide Synthesis Core and cloned into the CMV-SEAP plasmid to make CMV-Kozak-SEAP. The Kozak sequence with the α-granule targeting RANTES sequence was purchased as ultramers from Integrated DNA Technologies (IDT) and cloned into the CMV-SEAP to make the CMV-Kozak-RA-SEAP plasmid.

### Cell Transfections

For mouse MK cells, on day 10 of the differentiation process, cells in a single well of a 12-well plate were transfected with 1.6 μg of DNA using Lipofectamine 2000 (ThermoFisher Scientific #11668019) following the manufacturer’s instructions. The medium was changed 8-12 hours after transfection with fresh medium containing 10 ng/mL of recombinant mouse TPO. 48 hours after transfection MKs and PLPs were harvested for analysis. On day 12 of the differentiation process, the transfected cells were harvested for analysis.

For MEG-01 MK cells, 300,000 MEG-01 cells were initially plated in one well of a 24-well plate and transfected 24 hours later with 0.8 μg of DNA using Lipofectamine 2000 following the manufacturer’s instructions. For PLP loading, 500,000 MEG-01 cells were plated in a single well of a 6-well plate and 24 hours later were transfected with 4 μg of DNA using Lipofectamine 2000 following the manufacturer’s instructions. In both cases, the medium was changed 8-12 hours after transfection with fresh medium containing 100 ng/mL of recombinant human TPO. 48 hours after transfection MKs and PLPs were harvested for analysis.

### SEAP assay

SEAP levels were determined by adding QUANTI-Blue reagent (VWR # rep-qbs), whose change in color intensity from pink to purple/blue is proportional to the enzyme’s activity. In brief, medium was replaced 8-12 hours after transfection with fresh medium containing 10 ng/mL of TPO. For, ES derived platelets, on Day 12, PLTs loaded with SEAP were harvested and resuspended in 20 μL of OP9 media. Because FBS in the culture medium contains some basal alkaline phosphatase, all OD625 data were baseline corrected by subtracting the average basal OD625 value of the culture medium at each time point. Platelets were then activated with 1.92 U/mL of thrombin (Millipore Sigma) using a 50 U/mL stock solution that was prepared in PBS, 7.4 and 191.81 μM adenosine 5’-diphosphate (ADP; Millipore Sigma) that was from a stock solution of 25 mg/mL prepared in sterile Milli-Q water, before being added to the QUANTI-Blue reagent for time-course measurements. Separate wells were used for each time point in the 12-well platelet culture to maintain consistent conditions, varying only the incubation time. For each time point, 180 µL of QUANTI-Blue working solution was added to a 96-well plate, along with 20 µL of PLT suspension from SEAP-expressing platelets. The mixture was incubated at 37°C with 5% CO₂ for 24 hours, and SEAP activity was measured by recording the optical density at 625 nm (OD625). To determine the concentration of protein secreted by the cells, recombinant SEAP was serially diluted in culture medium at known concentrations and measured 24 hours later in assay media with an n=3 of independent replicates (Supplementary Fig. 14).

For MEG-01 derived PLPs, medium was replaced 8-12 hrs after transfection with fresh medium containing 100 ng/mL of TPO for ∼40 hr before starting the time-course measurements. For each time point, 180 μL of Quanti-Blue working stock was added to each well of a 96-well plate and 20 μL of cell culture supernatant from the SEAP-expressing cells and incubated at 37°C, 5% CO_2_ for 2 hrs. After incubation, SEAP activity was assessed by measuring the OD_625_ (Synergy HTX Reader, Biotek). Immediately after measuring the 3-hr timepoint, MKs and PLPs were activated with 1 U/mL of thrombin (Millipore Sigma) using a 50 U/mL stock solution that was prepared in PBS, 7.4 and 100 μM of adenosine 5’diphosphate (ADP; Millipore Sigma) that was from a stock solution of 25 mg/mL prepared in sterile Milli-Q water. Because FBS in the culture medium contains some basal alkaline phosphatase, all OD_625_ data were base-line corrected by subtracting the average basal OD_625_ value at each time point for untransfected MEG-01 cell culture medium. To assess the time for the reaction to reach a plateau, the CMV-SEAP plasmid was transfected in MEG-01 cells and the at OD_625_ was measured (Supplementary Fig. 15). Before base-line correction, we found that the enzymatic reaction reached a plateau at 2 hr. Baseline=corrected OD_625_ was plotted as a function of time using GraphPad Prism (GraphPad Inc). To determine the concentration of protein secreted by the cells, recombinant SEAP was serially diluted in culture medium at known concentrations and measured 2 hr later in assay media with an n=3 of independent replicates (Supplementary Fig. 15).

### Co-culturing mouse platelets and MEG-01 PLPs with HEK293-loxP-GFP-RFP (HEK293 reporter) cells

For mouse platelets, 10,000 HEK293 reporter cells were plated in one well of a 12-well plate and allowed to recover for 48 hours. Platelets loaded with Cre-ER were isolated from two wells of a 12-well plate and resuspended in HEK293 growth medium. The loaded platelets were transferred to one well of a 12 well-plate containing the HEK293 reporter cells. To facilitate the translocation of Cre recombinase to the nucleus of the HEK293 reporter cells, a 20 mg/mL stock solution of 4-OHT (Millipore Sigma #H7904) was prepared in ethanol and diluted to 1mg/mL in PBS and added to the medium to a final concentration of 4 ng/mL. 48 hours after co-incubation, the HEK293 reporter cells were analyzed by flow cytometry.

For MEG-01 PLPs, 10,000 HEK293 reporter cells were plated in one well of a 12-well plate. The next day PLPs loaded with Cre-ER from 2 wells of transfected MEG-01 were collected and resuspended in HEK293 growth medium. The loaded PLPs were transferred to one well of a 12-well plate containing the HEK293 reporter cells. To enable the Cre recombinase to translocate to the nucleus of the HEK293 reporter cells, a 1 mg/mL stock solution of 4-OHT (Millipore Sigma #H7904) was prepared in ethanol and added to the medium to a final concentration of 40 ng/mL. 48 hours after co-incubation, the HEK293 reporter cells were analyzed by flow cytometry.

### Preparation for flow cytometry

Once the cells were collected, they were resuspended in 100 μL PBS containing 3% BSA for antibody labeling. For nucleated cells (MEG-01, MK, and HEK293 reporter line), cells were stained with 1:200 dilution of 1 mg/mL stock of Hoechst 33342 and incubated at 37°C, 5% CO_2_ for 30 minutes. After incubation, the cells were washed with PBS and resuspended in 500 μL of PBS for flow cytometry analysis. The anucleated cells (mouse platelets and MEG-01 PLPs) were stained with calcein red-orange AM (ThermoFisher #C34851) at a dilution of 1:200 from a 1mM stock following the manufacture’s protocol. For confirming mESC derived mature MKs and platelets, FITC-conjugated anti-mouse CD41 (Biolegend 133903) was used at 1:200 dilution. Platelet activation was assessed with APC anti-mouse P-selectin/CD62P (Fisher, 148303) that was used at 1:200. For confirming MEG-01 derived mature MKs and PLPs, APC-conjugated anti-human CD41 (Fisher #559777) was diluted at 1:100.

### Flow cytometry

Cells were washed and resuspended in PBS. Fluorescence data were collected using a Beckman Coulter Life Sciences CytoFLEX S cytometer. Alexa Fluor 488 and FITC was measured using a 488 laser and a 530/30 filter. APC and Alexa Fluor 647 were measured using a 640 laser with 660/10 filter. All flow cytometry data was analyzed using FlowJo.

### Preparation for microscopy

For the analysis of GFP and VWF expression using microscopy, coverslips (Fisher Scientific #12541005) were prepared by UV sterilizing each side with UV light for 30 minutes. After sterilization, the coverslips were placed in a 12-well plate and a few drops of poly-D-lysine (PDL) (VWR #A3890401) were added onto each coverslip. The plate with coverslips was incubated overnight in a humidified 5% CO_2_, 37 °C to allow the PDL to coat the coverslips. The following day, the PDL was aspirated off the coverslips and left to dry in a humidified 5% CO_2_, 37 °C incubator overnight. On day 7 of mESC differentiation, 30,000 OP9 cells were plated onto the treated coverslips. The next day, day 8 of mESC differentiation, the differentiating mESCs were treated with 0.25% trypsin to detach cells from the well, neutralized with MEF growth medium, centrifuged at 300 xg for 5 minutes and resuspended in OP9 medium with 10 ng/mL TPO then added to the plated OP9 cells. Cells were transfected with respective plasmids on day 10 of differentiation and a media swap containing 10 ng/mL of TPO was done 6-8 hours after transfection. On day 12, cells were fixed with 4% paraformaldehyde (VWR #100503-917) for 10 minutes at room temperature, washed with PBS and permeabilized using a blocking solution (5% donkey serum, Jackson ImmunoResearch #017-000-121 with 1% BSA, Jackson ImmunoResearch #001-000-162) supplemented with 0.25% Triton-X100 for 10 minutes at room temperature and washed with PBS. For a complete description of staining reagents, see Supplementary Table S1.

### Microscopy

Images that were taken from a slide (Figs. 1b, 2b, and 2k) were acquired using a Zeiss LSM800 confocal microscope with a 100x objective. Images were processed using Zeiss Zen software and figures were prepared using Adobe Illustrator. Images that were taken from a tissue culture plate (Figs. 2f and g) were acquired using a Nikon TS100 microscope equipped with a SOLA light engine, 380-680 nm wavelength range and a Nikon EF-4 Endow GFP HYQ excitation/emission filter, and a ProgRes high resolution monochrome camera. In all imaging, care was taken to not oversaturate the fluorescence measurements.

### Online content

Methods, including statements of data availability and any associated accession codes and references are available at (link inserted by journal).

## Data Availability

All data collected to evaluate the conclusions of this work are presented in the paper and/or in the Supplementary Materials. All plasmids generated from this study will be deposited in Addgene after publication. Reagents or additional data are available from the corresponding author upon request.

## Acknowledgements

We thank J. Marvin at the University of Utah Flow Cytometry Core for his assistance with our flow cytometry experiments and support from their Core Flow Cytometry grants from the Office of the Director of the National Institutes of Health under Award Number S10OD026959 and NCI Award Number 5P30CA042014-24. We also thank the DNA/Peptide Synthesis Core at the University of Utah for oligonucleotide synthesis. This work was supported by the National Institute of Health Trailblazer Award 1R21EB025413-01, awarded to TLD, the National Institute of Health Director New Innovator Award 1DP2CA250006-01, awarded to TLD, and National Institute of Health F31 1F31HL167592-01 awarded to SJ. We would also like to thank Lydia Taylor for critically reading and evaluating the manuscript.

## Author contributions

TLD conceived of the study and designed the experiments. FI conducted MEG-01 transfections with GFP, and all the mESC differentiation studies with mouse platelets. FI also conducted all of the scale-up experiments in both mESCs and MEG-01 cells. SBJ constructed the α-granule targeted GFP plasmid and the SEAP plasmid used in the MEG-01 cells. SJ also carried out the Cre and SEAP experiments in MEG-01 cells. TLD constructed the Cre plasmid and performed all the microscopy in this study. MRL and JC built the Kozak-SEAP plasmids used for the SEAP experiments in mESCs, helped with cell maintenance, and transfections. HW helped with cell maintenance and transfections. TLD and FI wrote the manuscript with contributions from SBJ, MRL, JDC, and HW.

## Additional information

Supplementary information is available for this paper at (link inserted by journal).

## SUPPLEMENTARY INFORMATION

**Supplementary Figure 1.**
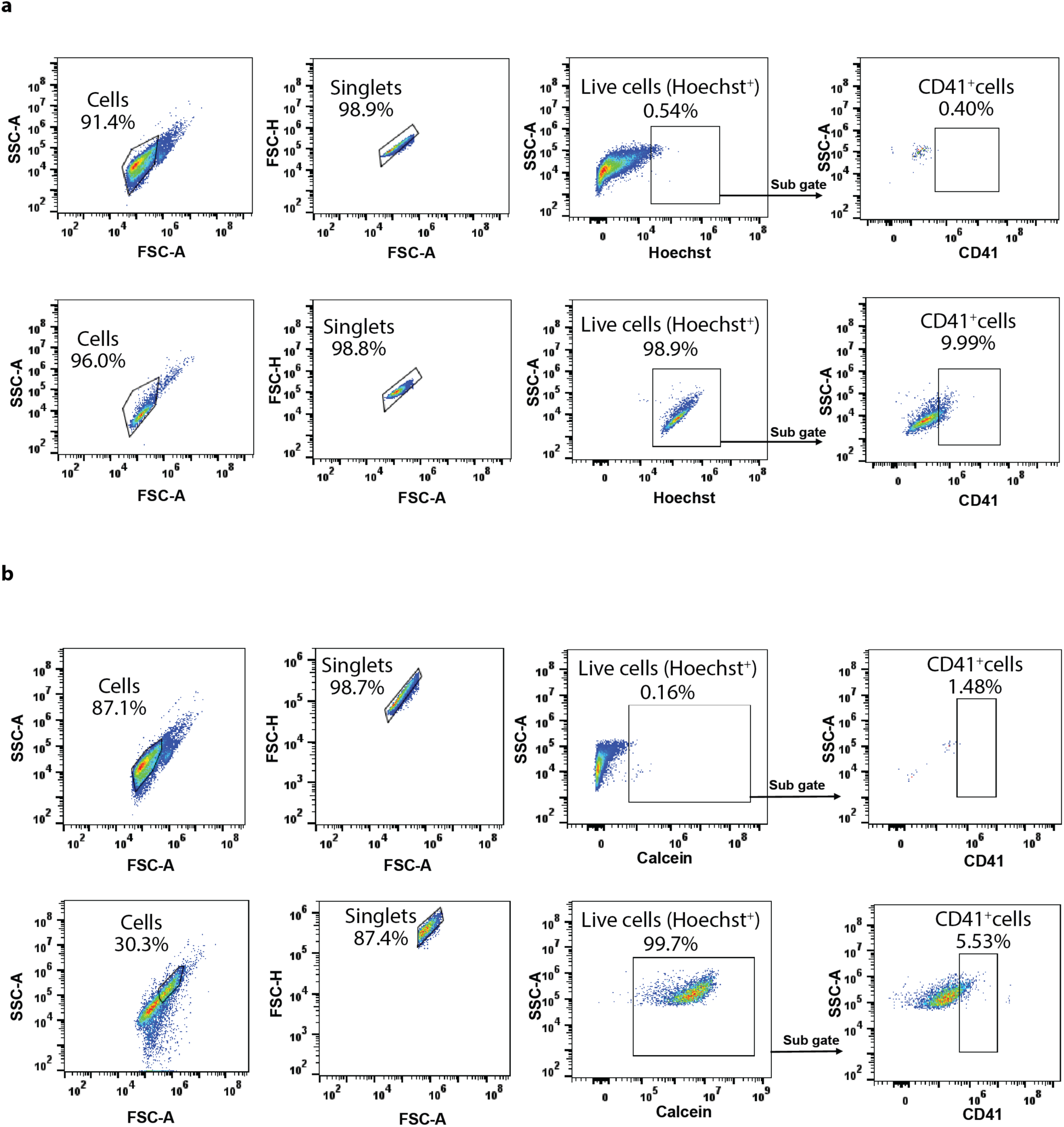
Gating strategy for MKs and platelets. **a** For defining areas for the gating strategy, non-labeled cells were observed in the flow cytometer (top row). Observing CD41+ MKs, single cells (e.g. singlets) were identified and the live cells were gated from this population as Hoechst^+^ cells and analyzed for CD41^+^ staining (bottom row). **b** A similar gating strategy was used for identifying CD41+ platelets with defining the unlabeled cells (top row) and labeled cells (bottom row). Here, calcein staining was used to identify the live cells since platelets are anucleate.

**Supplementary Figure 2.**
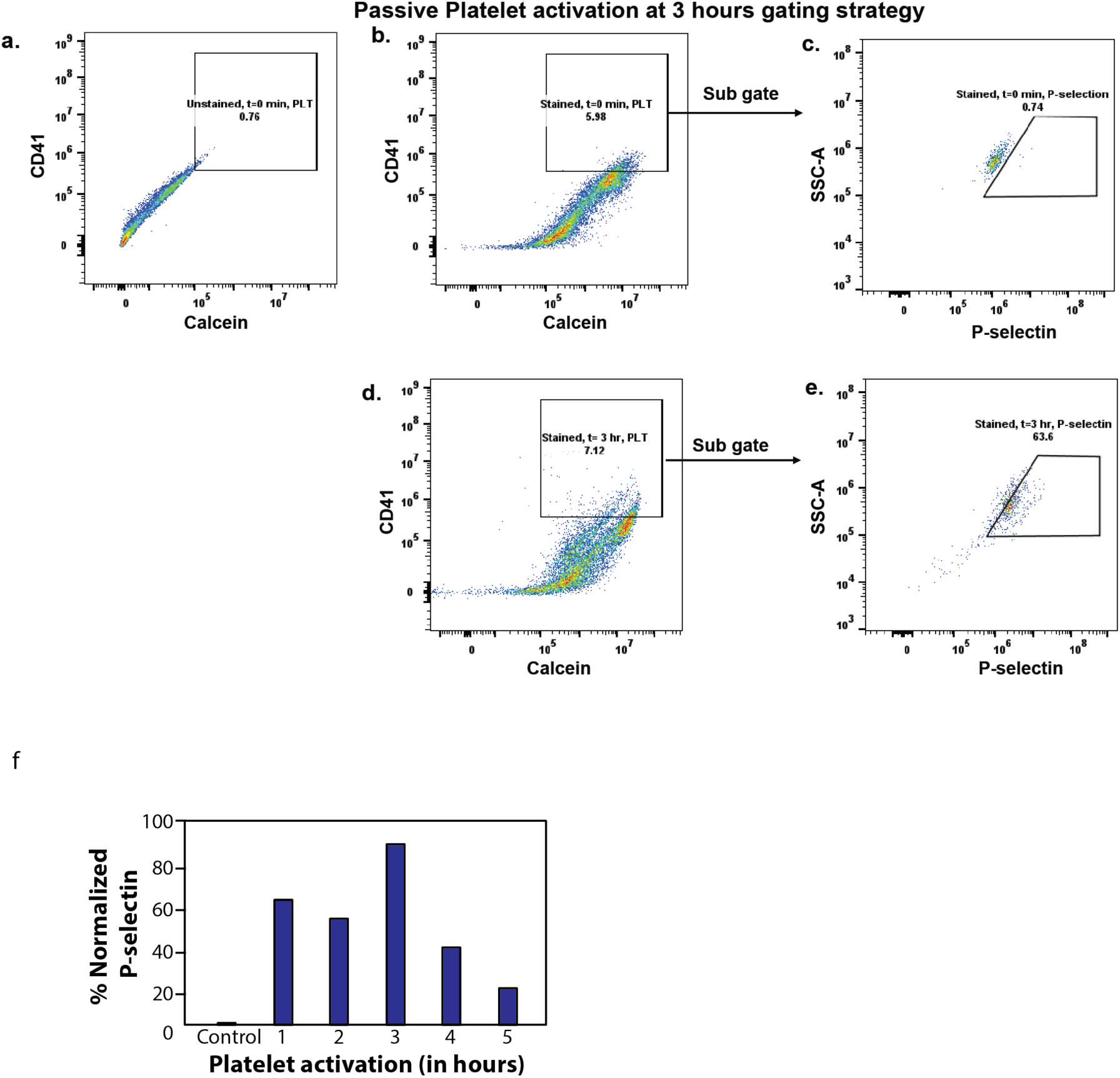
Activation of mouse platelets. **a** Unstained negative control for identifying boundaries for calcein+ cells and CD41+ expression. **b** Stained platelets with calcein and CD41 at t=0 minutes with a gate drawn to determine the platelet population based on negative control staining. **c** The platelet population from b was analyzed for P-selectin expression at t=0 minutes. **d** Platelets were identified after 3 hours of exposure to thrombin **e** P-selectin expression at t=3 hours. **f** Percentage of P-selectin expression in platelets at various time points after activation. The control represents p-selectin expression at time t=0, immediately after the addition of thrombin/ADP. P-selectin expression at subsequent time points (t=1, 2, 3, 4, and 5 hours) has been normalized to this control. The percentage indicates the proportion of platelets expressing p-selectin at each time point.

**Supplementary Fig. 3:**
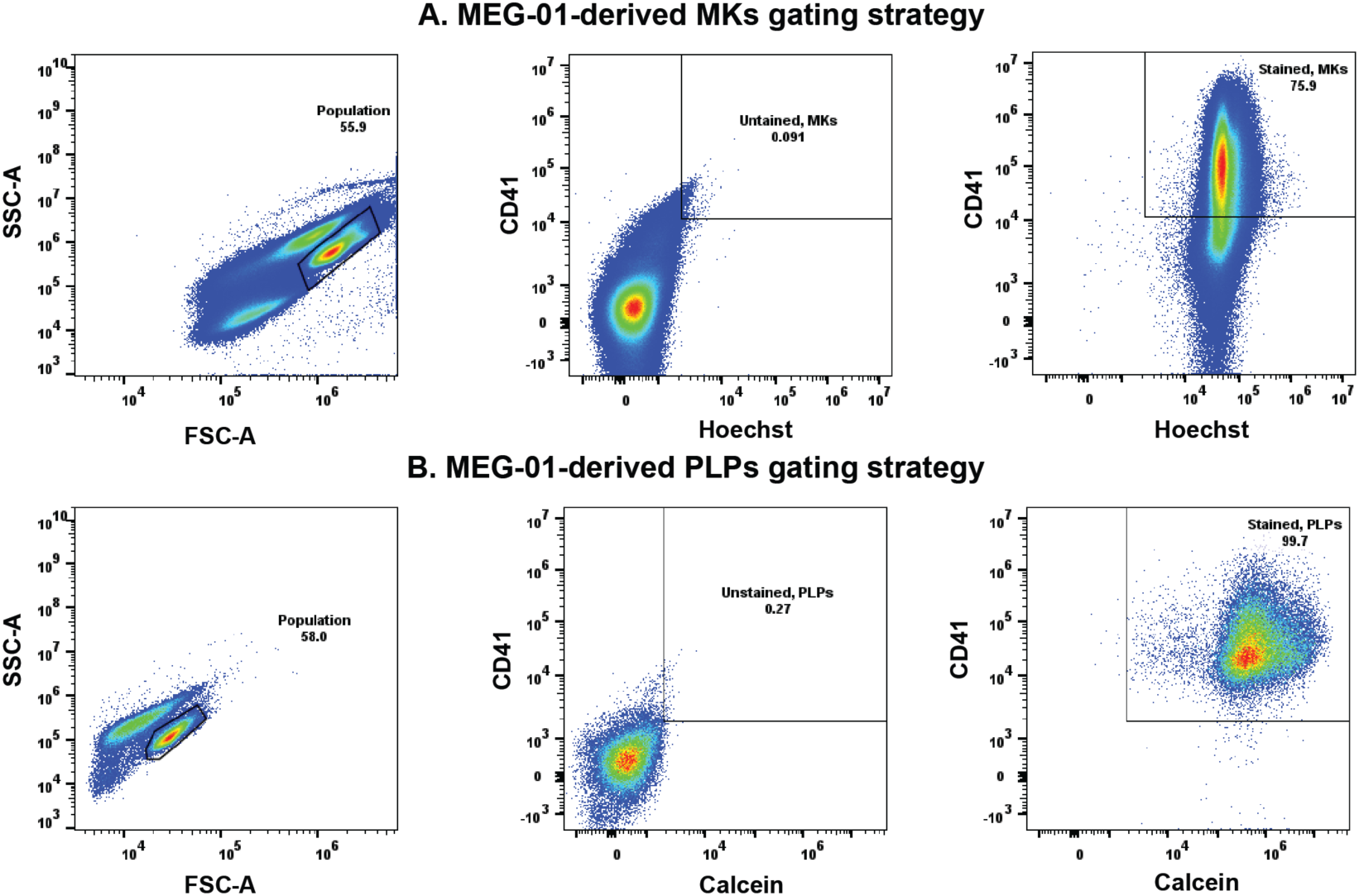
Gating strategy for MEG-01 cells and PLPs. **a** MEG-01 cells were identified based on forward scatter area (FSC-A) and side scatter area (SSC-A) (left). Negative control of MEG-01 cells not stained identified the gating strategy for the Hoechst^+^ and CD41^+^ cells (middle), which identifies where the mature MKs derived from MEG-01s are seen after staining (far right). b PLPs were identified based on forward scatter area (FSC-A) and side scatter area (SSC-A) (left). Negative control of PLPs not stained with CD41 or calcein (middle), which indicates the PLP population after staining (far right). Each population was gated separately and analyzed for Hoechst/calcein and CD41 markers. Populations negative for Hoechst and calcein were excluded and CD41^-^ indicated immature MKs or PLPs.

**Supplementary Figure 4:**
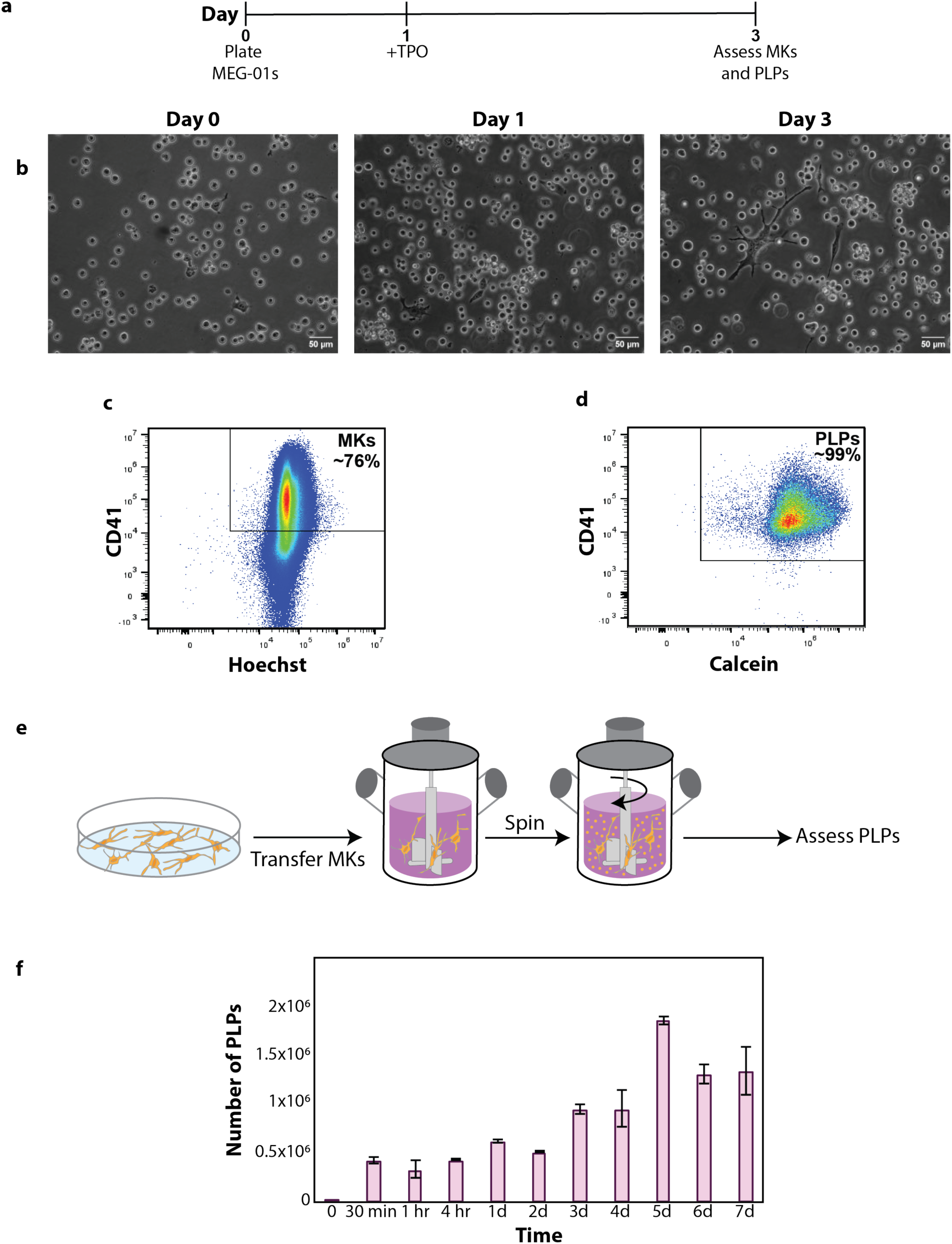
Characterization of MEG-01 maturation and platelet-like particles (PLP) production. **a** Timeline of MEG-01 maturation in the presence of TPO. **b** Representative images of MEG-01 cell maturation. **c** Flow cytometry analysis of MKs after three days of maturation in TPO with a box drawn to represent CD41^+^ and Hoechst^+^ cells. **d** Flow cytometry analysis of PLPs after being harvested from growing MEG-01s in the presence of TPO for 3 days with a box drawn to represent CD41^+^ and Calcein^+^ cells. **e** MEG-01s were matured in the presence of TPO for 3 days and the entire culture was collected, transferred to a spinner flask bioreactor and analyzed at various time points. **f** PLP production in spinner flask bioreactors were analyzed at various time points after MKs were added. The time ranged from minutes (min) to hours (hr) and days (d). The specified number of PLPs represents the average of 2 biological replicates. The error bars represent the standard deviation of the mean.

**Supplementary Figure 5:**
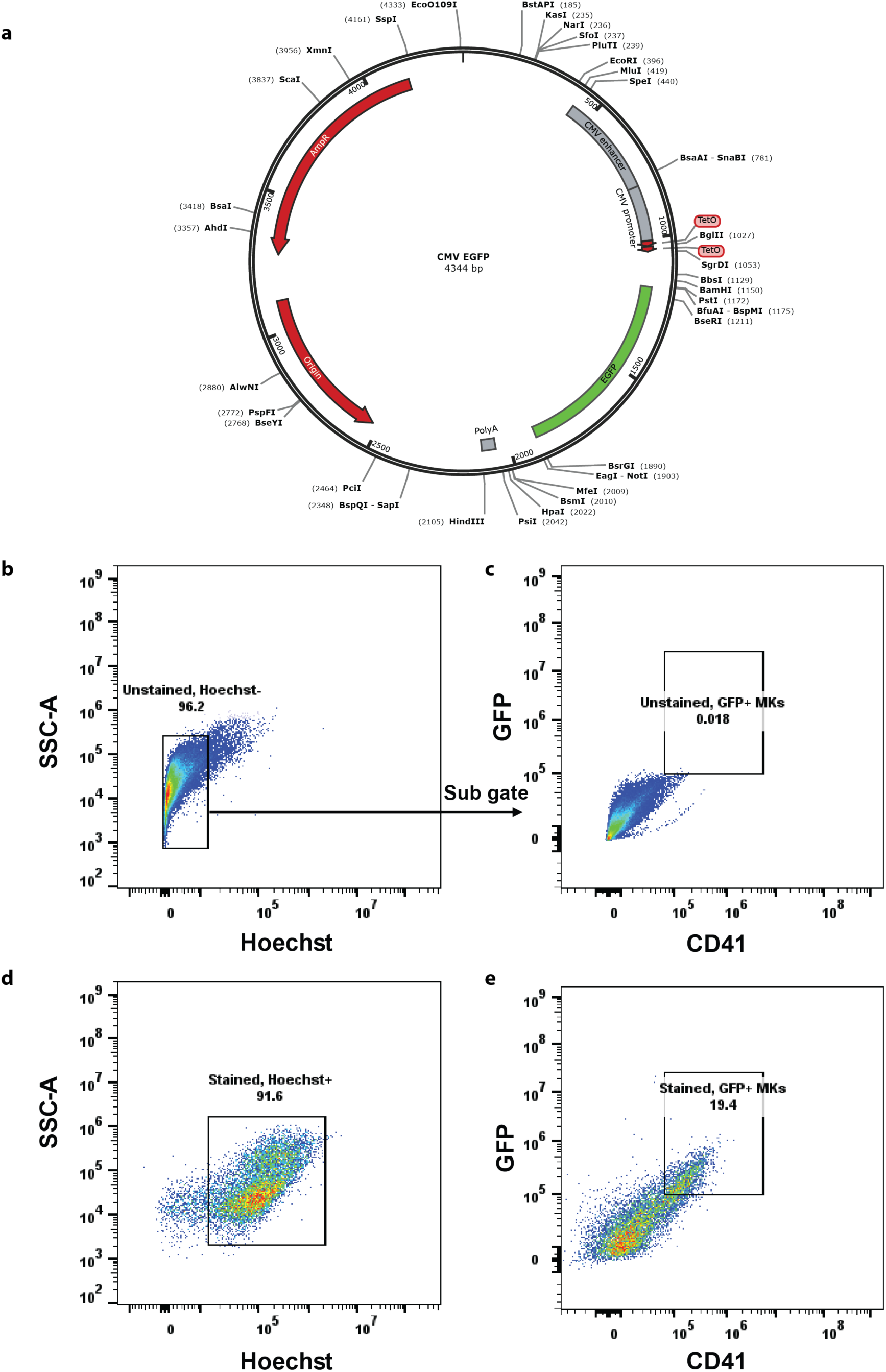
Gating strategy for GFP+ MKs. **a** mESCs were differentiated to MKs and on day 10 of differentiation, MKs were transfected with a plasmid that constitutively expresses GFP. **b** Flow cytometry of non-labeled cells as the negative control to identify gating parameters. **c** The Hoechst-negative population was gated to identify GFP^-^ and CD41^-^ regions. **d** GFP transfected MKs stained with Hoechst. The Hoechst+ cells were gated and **e** GFP and CD41 labeling were assessed.

**Supplementary Figure 6.**
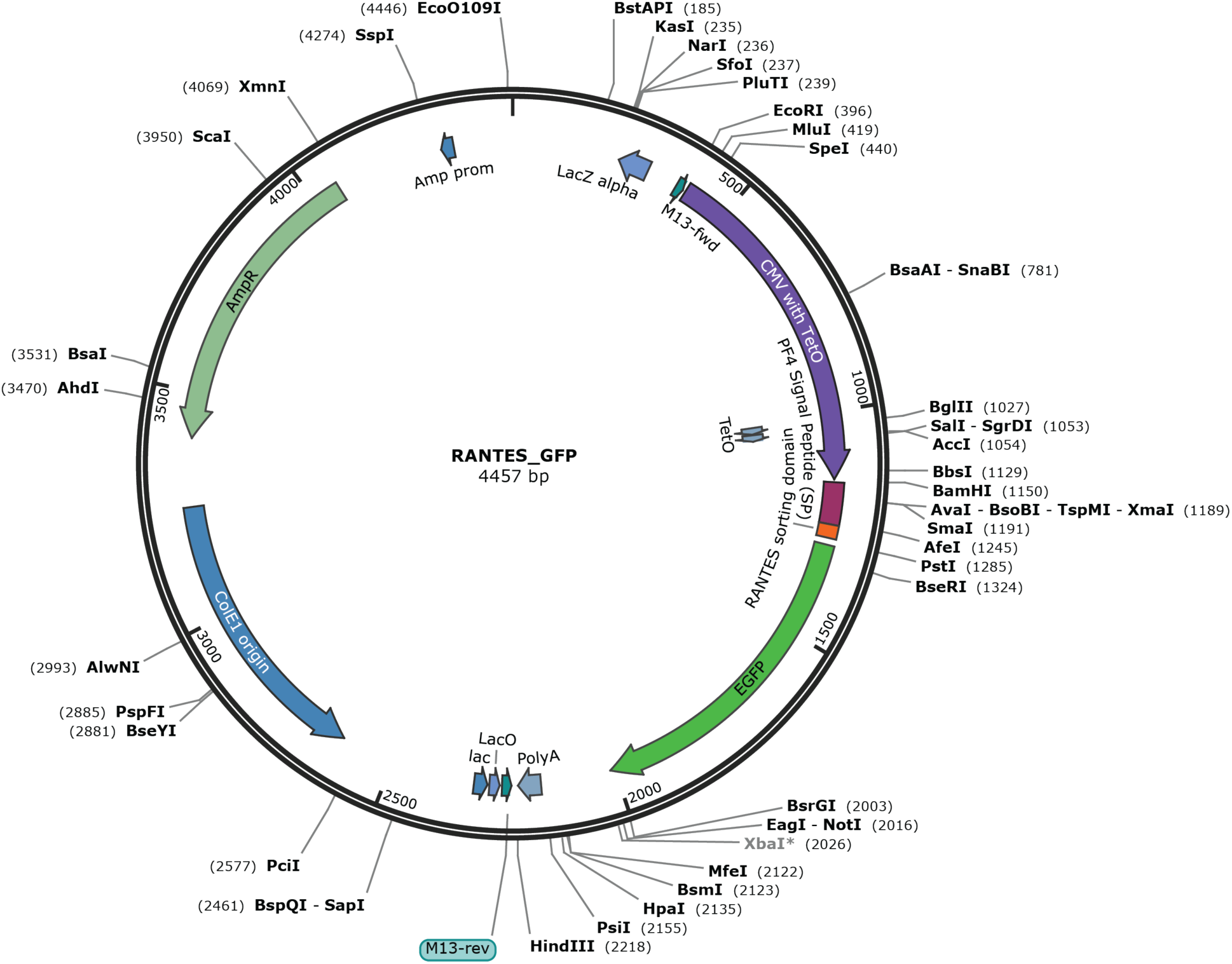
RANTES GFP plasmid. Plasmid constitutively expressing GFP fused to peptides for targeting GFP to α-granules.

**Supplementary Figure 7:**
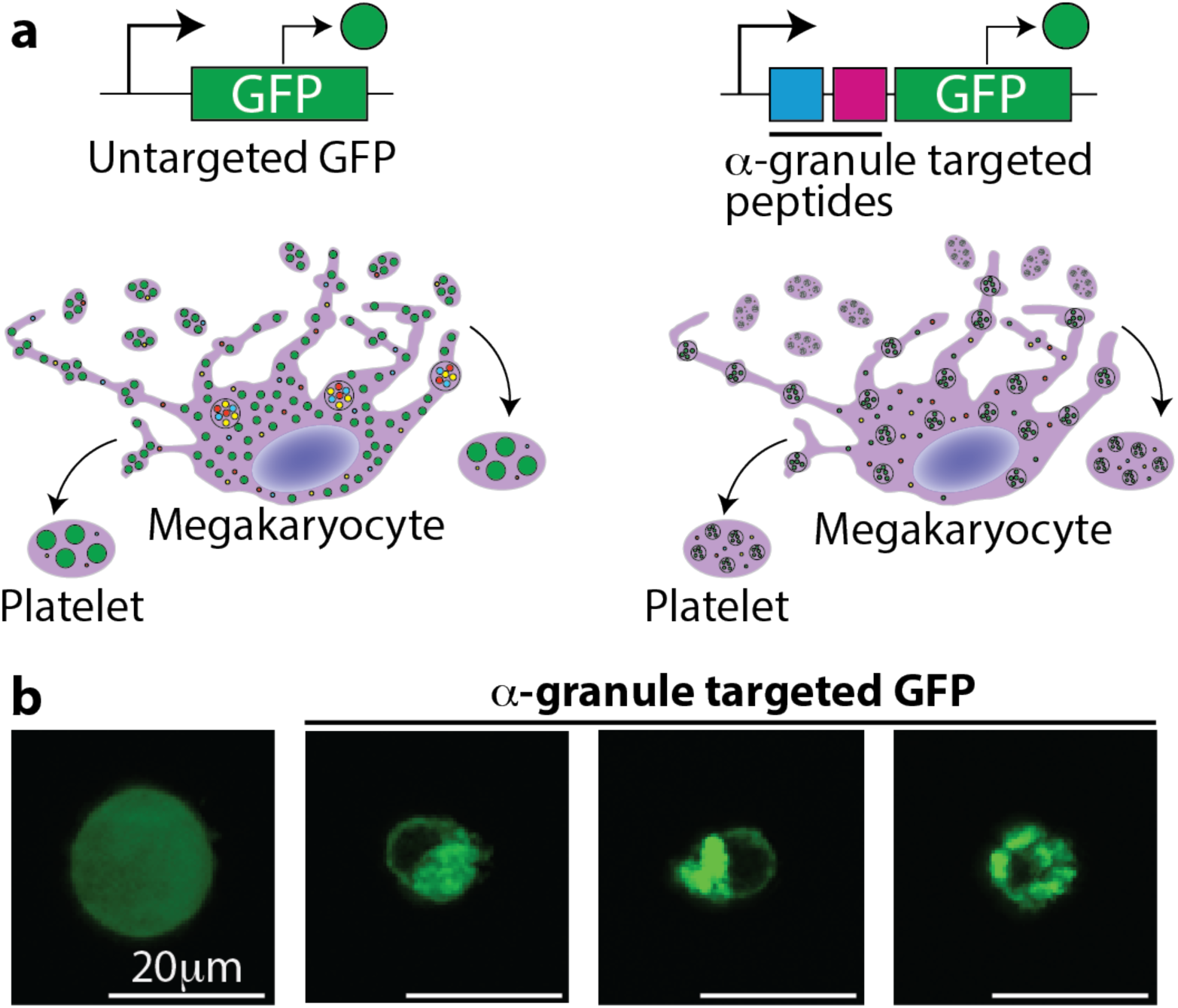
Targeting proteins to α-granules in human MEG-01 cells. **a** Schematic of non-α-granule targeted GFP proteins in MEG-01s (left) and GFP fused to peptides for targeting to α-granules (right). **b** Fluorescent image of non-targeted GFP in MEG-01s (left) compared to GFP that is targeted to the α-granules (right).

**Supplementary Figure 8.**
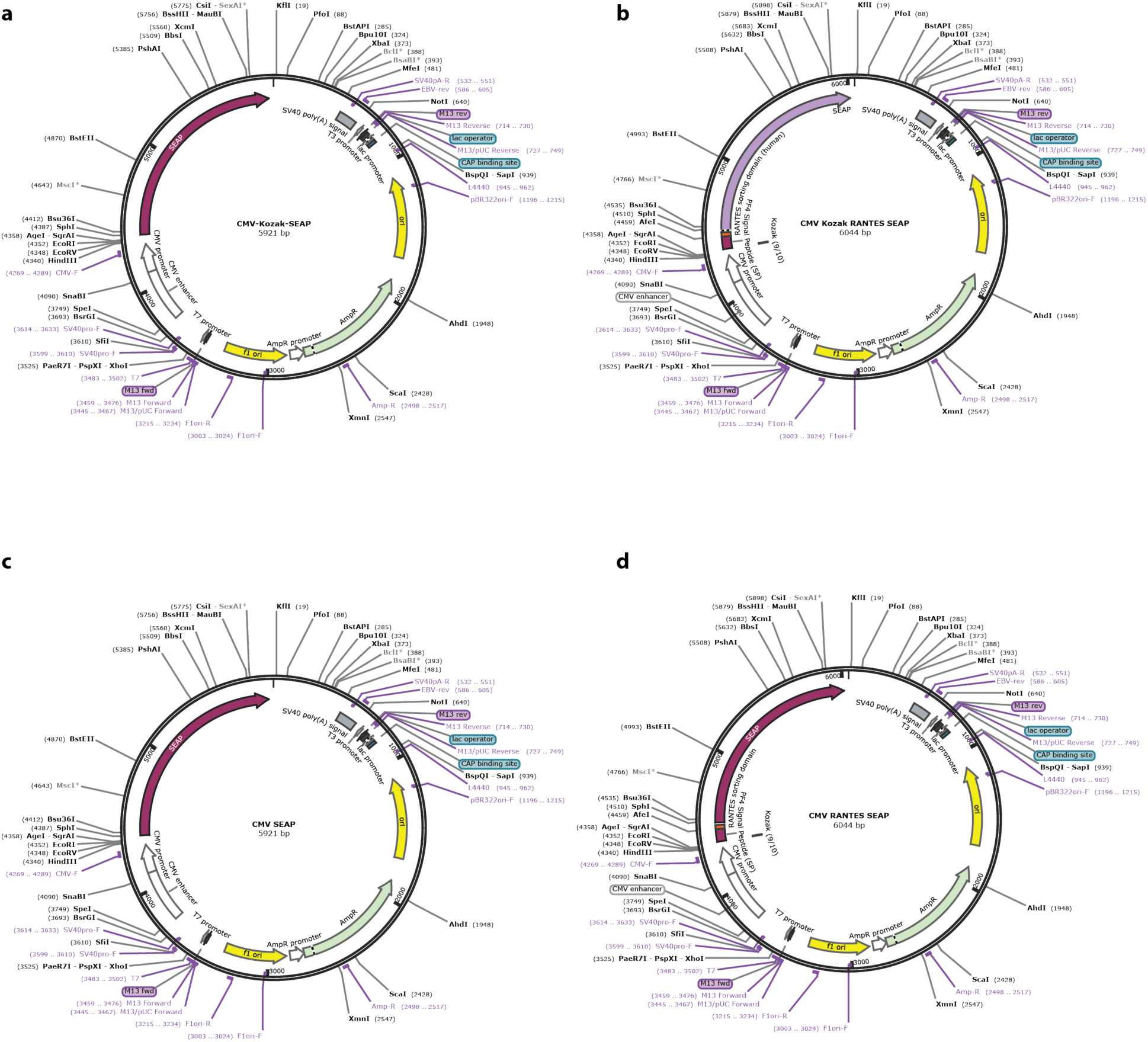
SEAP plasmids. **a** Plasmid constitutively expressing SEAP used in mouse MKs and platelets. **b** Plasmid constitutively expression SEAP fused to a short peptide sorting signal derived from the human cytokine RANTES used in mouse MKs and platelets to target SEAP to the α-granules. **c** Plasmid constitutively expressing SEAP used in human MEG-01 MKs and PLPs. **d** Plasmid constitutively expression SEAP fused to a short peptide sorting signal derived from the human cytokine RANTES used in human MEG-01 MKs and PLPs to target SEAP to the α-granules.

**Supplementary Figure 9.**
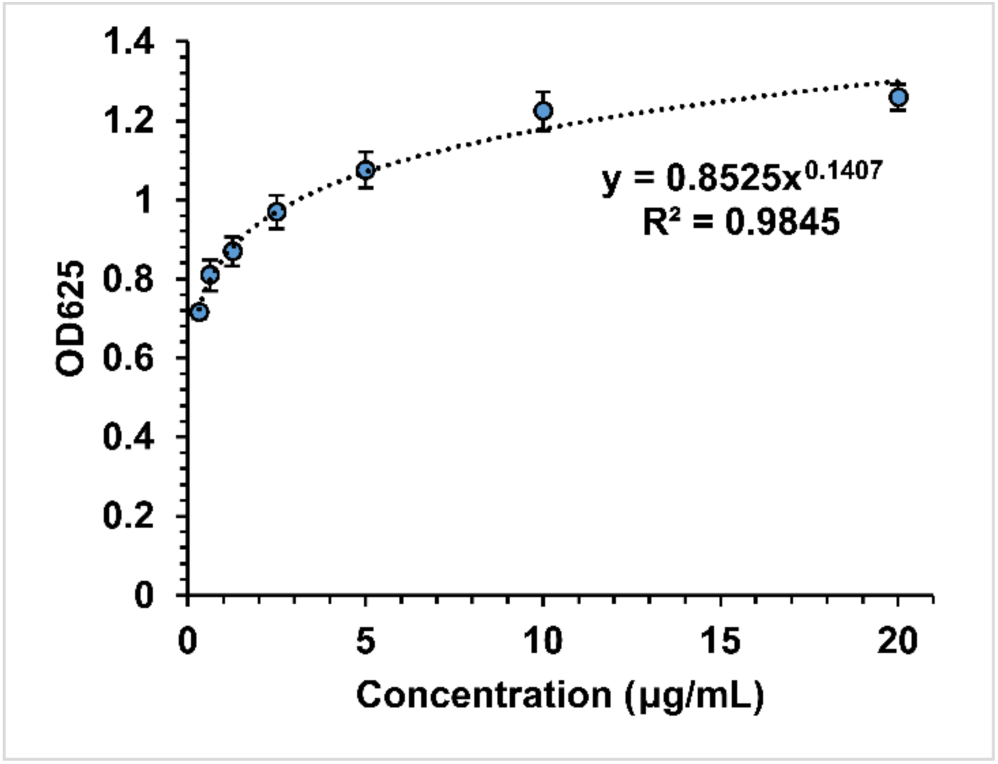
SEAP calibration curve for converting OD_625_ values to concentration. The optical density (OD_625_) of recombinant human SEAP serially diluted in OP9 culture media at known concentrations was measured after 24 hours in assay media (n = 3 independent replicates ± SD). Microsoft Excel was used to fit the data to a power trendline for converting experimental OD_625_ values into concentration values of SEAP.

**Supplementary Figure 10.**
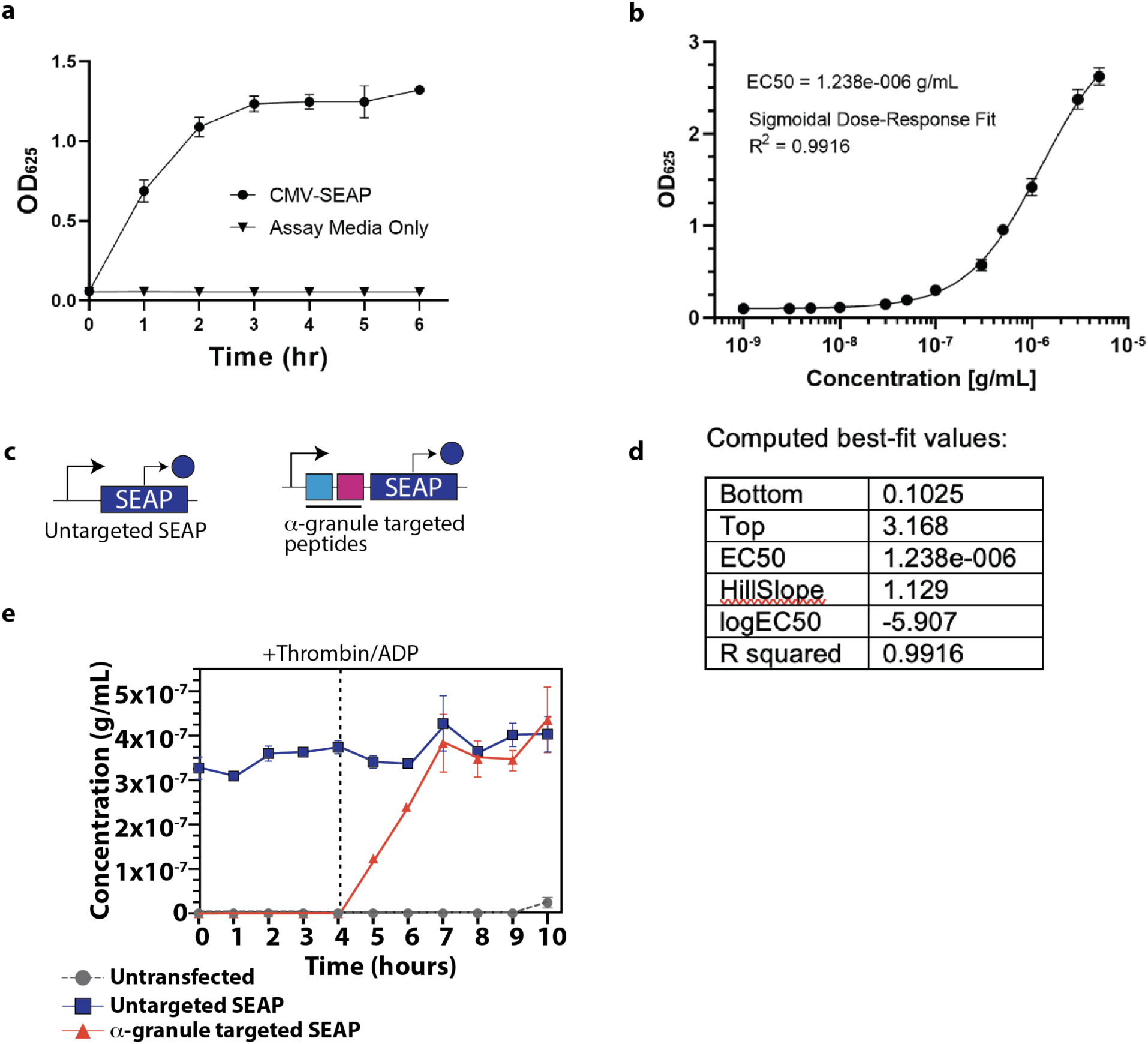
SEAP calibration curve for converting OD_625_ values to concentration medium and release of SEAP in MEG-01. **a** Kinetics of QUANTI-Blue colorimetric assay with CMV-SEAP plasmid in MEG-01 medium. Non-α-granule-targeted SEAP (black circles) vs. time. These experiments were repeated independently at least three times with similar results. Error bars indicate the standard error of the mean. **b** SEAP calibration curve for converting OD_625_ values to concentration. The optical density (OD_625_) of recombinant human SEAP serially diluted in culture media at known concentrations was measured after 2 hours in assay media (n = 3 independent replicates ± SEM). **c** Schematic of constitutively expressing SEAP (left) and SEAP fused to a short peptide sorting signal derived from the human cytokine RANTES used in human MEG-01 MKs and PLPs to target SEAP to the α-granules. **d** GraphPad prism was used to fit the data to a sigmoidal dose-response curve for converting experimental OD_625_ values into concentration values of SEAP, using the standard model of: 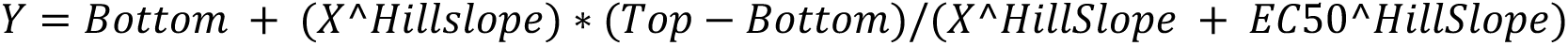 **e** Non-α-granules targeted SEAP and its release from PLPs over time (blue squares). SEAP targeted to the α-granule (red triangles) and its release upon activation with the addition of thrombin/ADP (dotted line). These experiments were repeated independently at least three times with similar results. Error bars indicate the standard error of the mean.

**Supplementary Figure 11.**
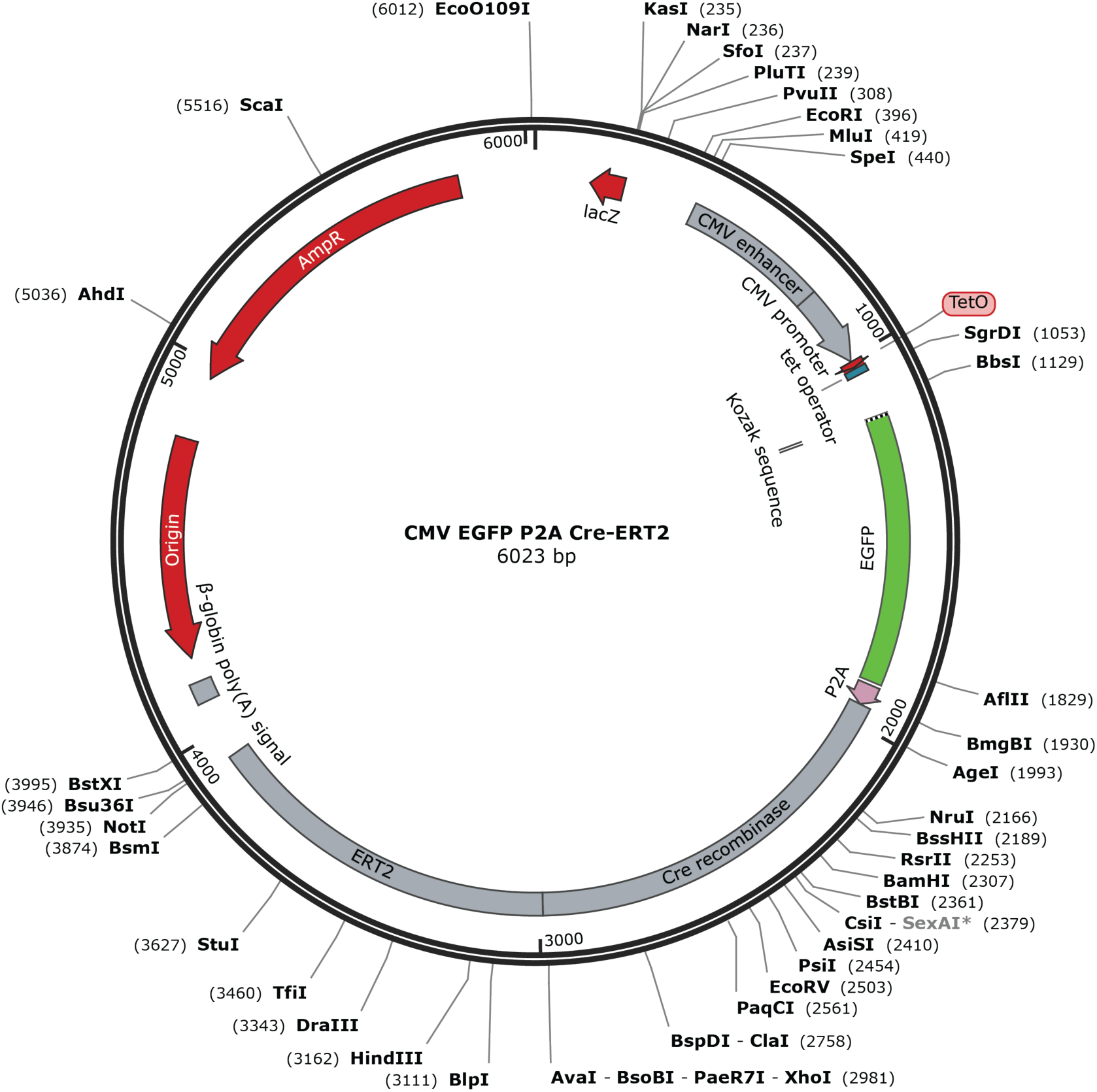
GFP-P2A-Cre plasmid. GFP is constitutively expressed along with Cre-ER.

**Supplementary Figure 12:**
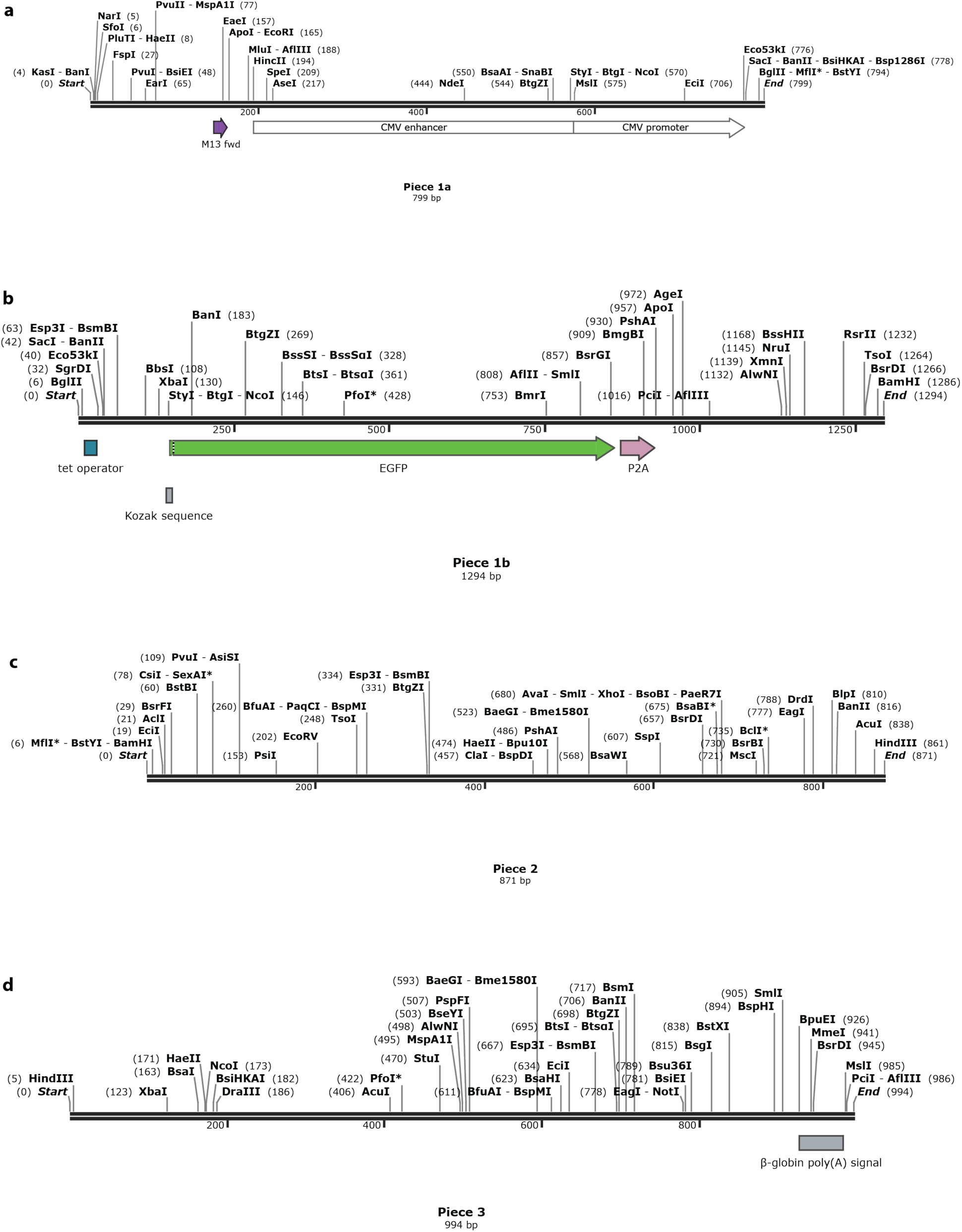
Building the GFP-P2A-Cre plasmid.

**Supplementary Figure 13:**
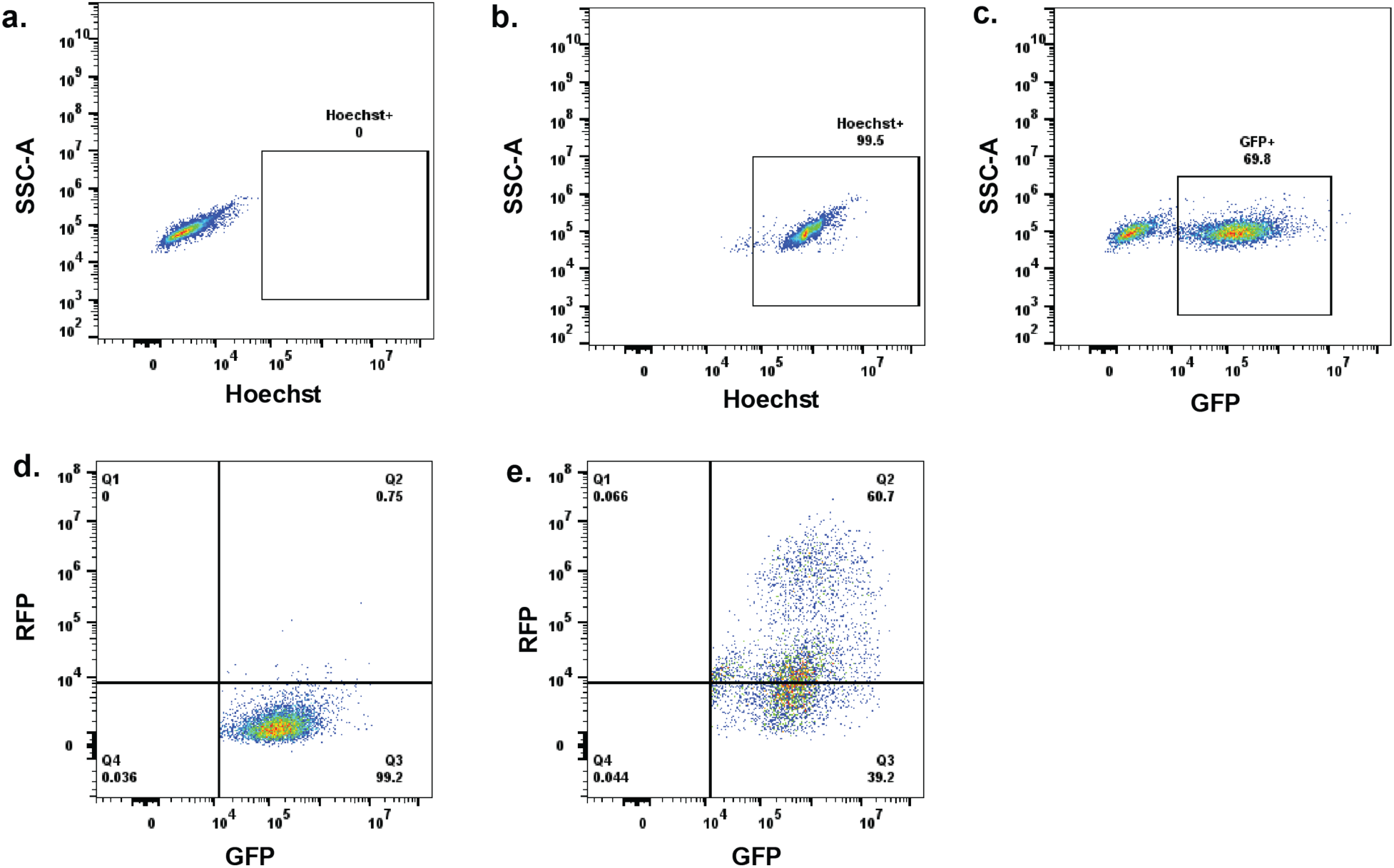
HEK293 reporter cell line gating strategy. **a** HEK293-loxP-GFP-RFP cells in the absence of co-cultured engineered platelets (untreated) and not stained with Hoechst. **b** HEK293-loxP-GFP-RFP cells stained with Hoechst and in the absence of co-cultured engineered platelets (untreated). **c** Hoechst^+^ cells (live cells) were gated and GFP expression of these cells was assessed. The GFP^+^ cells were gated since these cells have the functional reporter. **d** Untreated GFP^+^ cells and RFP^-^ cells. **e** 48 hours after co-culturing engineered platelets filled with Cre and 4-OHT was added to the media using the gating strategy described above.

**Supplementary Fig. 14.**
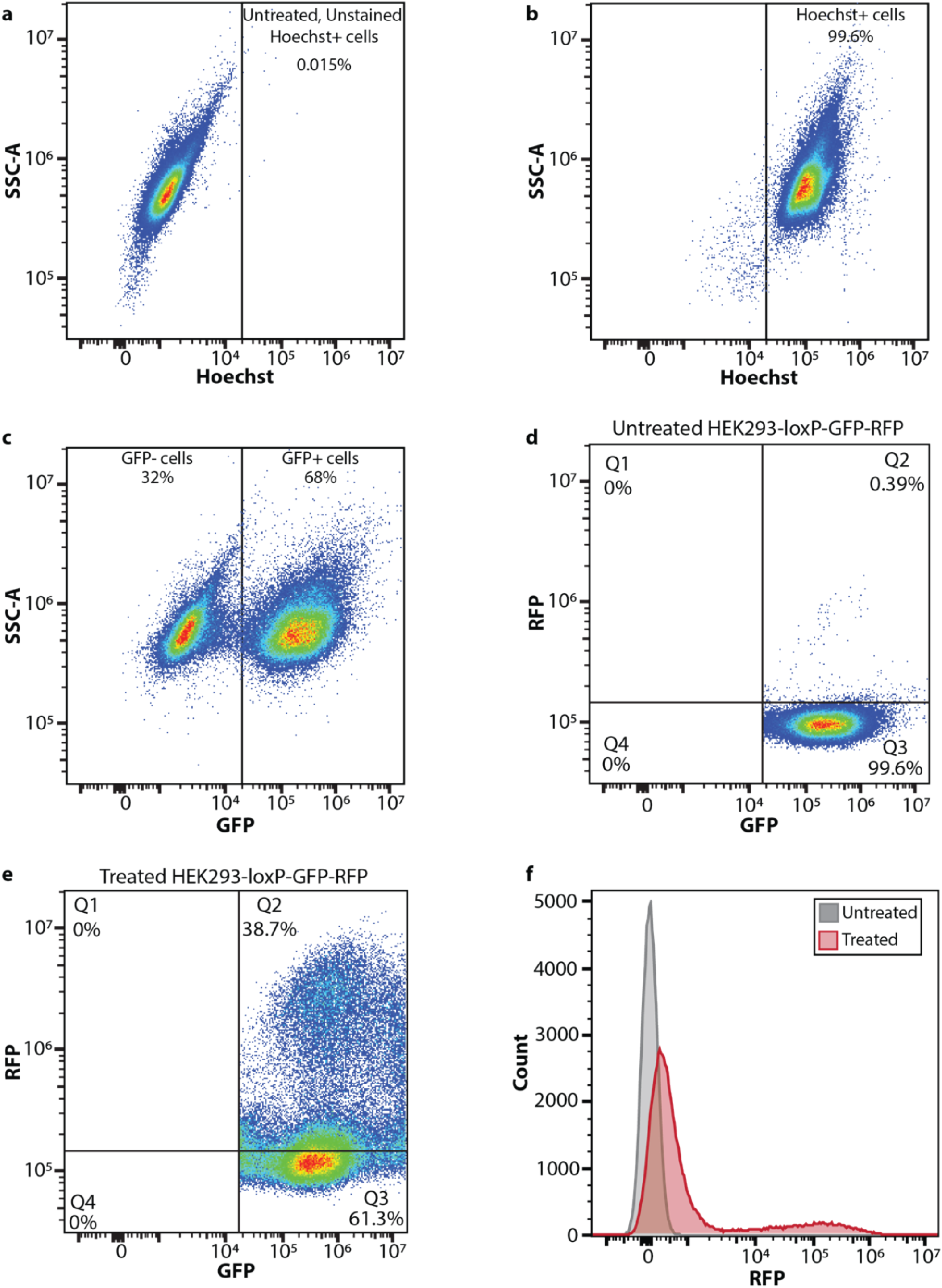
Flow cytometry gating strategy for HEK293-loxP-GFP-RFP analysis. **a** HEK293-loxP-GFP-RFP cells in the absence of co-cultured engineered PLPs (untreated) and not stained with Hoechst. **b** HEK293-loxP-GFP-RFP cells in the absence of co-cultured engineered PLPs and stained with Hoechst to identify the live cells (right rectangle). **c** The live cells were gated and analyzed for GFP expression (right rectangle) in the absence of co-cultured engineered PLPs. **d** The untreated GFP positive cells were analyzed for RFP expression (Q2). **e** After co-culturing engineered PLPs filled with Cre/ER and 4-OHT for 48 hrs, the HEK293-loxP-GFP-RFP cells were gated the same way as described for the untreated cells and RFP expression was quantified (Q2). **f** Histogram comparing the RFP expression of untreated (grey) and treated (red) HEK293-loxP-GFP-RFP cells.

Key Resource.

| Key Resource Table S1 |  |  |  |  |
| --- | --- | --- | --- | --- |
| Resource reagent | Description | Source | Catalog # | Additional Information |
| Primary antibody | Rabbit anti Von Willebrand factor (VWF) | Fisher Scientific | NB60058601M | Dilution used: 1:200 |
| Primary antibody | Goat anti-GFP | ThermoFisher | 600-101-215M | Dilution used: 1:500 |
| Conjugated antibody | APC anti-mouse/rat CD62P (P-selectin) | Fisher Scientific | 148303 | Dilution used: 1:200 |
| Conjugated antibody | FITC-conjugated anti-mouse CD41 | Biolegend | 133903 | Dilution used: 1:200 |
| Conjugated antibody | APC-conjugated anti-human CD41 | Fisher | 559777 | Dilution used: 1:100 |
| Secondary antibody | Alexa Fluor 488 AffiniPure Donkey anti goat IgG (H+L) | Jackson ImmunoResearch | 705-545-147 | Dilution used: 1:1500 |
| Secondary antibody | Alexa Fluor 647 AffiniPure Donkey anti Rabbit IgG (H+L) | Jackson ImmunoResearch | 711-605-152 | Dilution used: 1:1500 |
| DNA dye | Hoechst 33342 | BD Pharmingen | 551908 | Dilution used: 1:200 |
| Cell viability dye | Calcein red-orange AM | ThermoFisher | C34851 | Dilution used: 1:200 |
| Software | FlowJo® | FlowJo, LLC | NA |  |

**Table S1.**
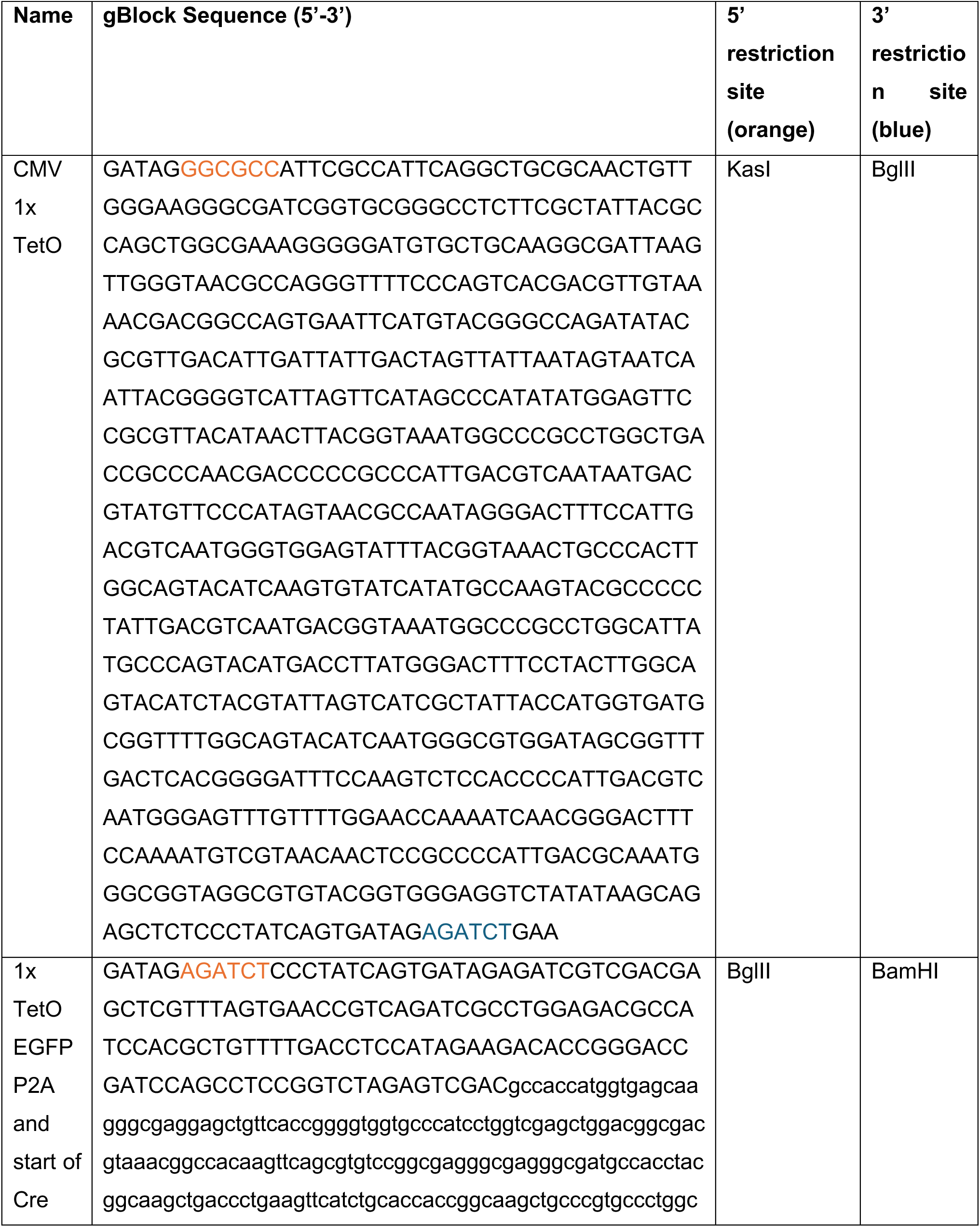
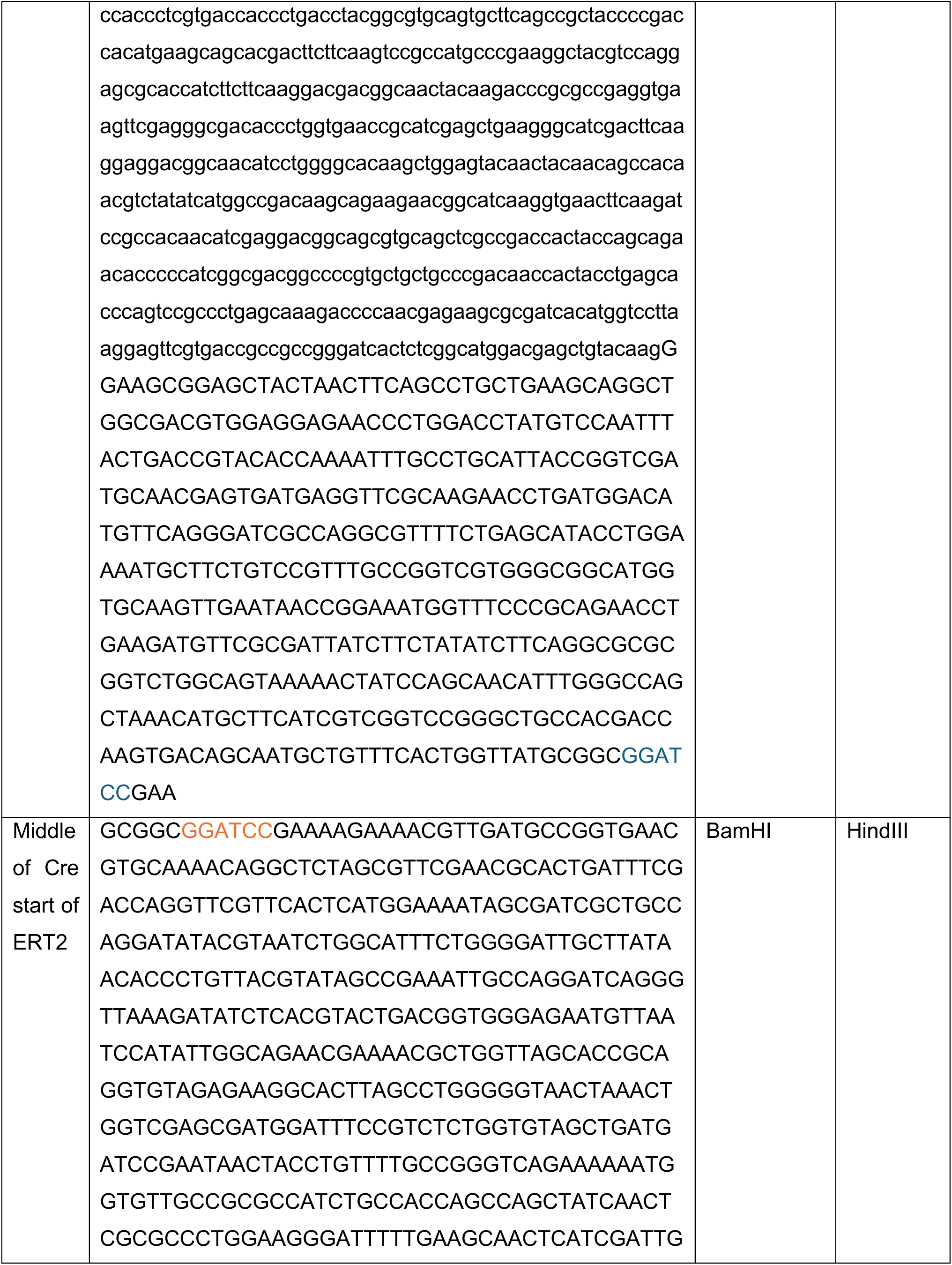
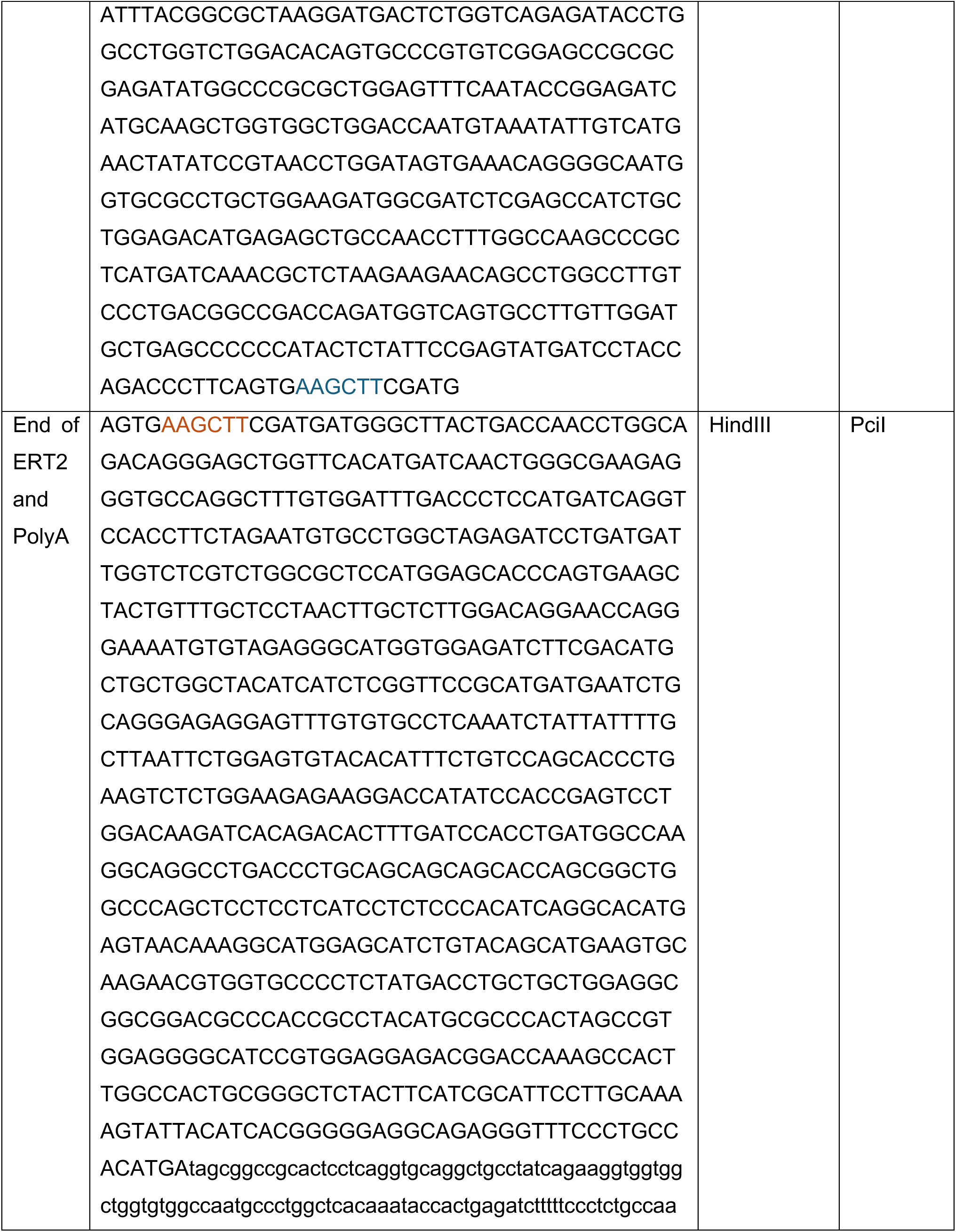
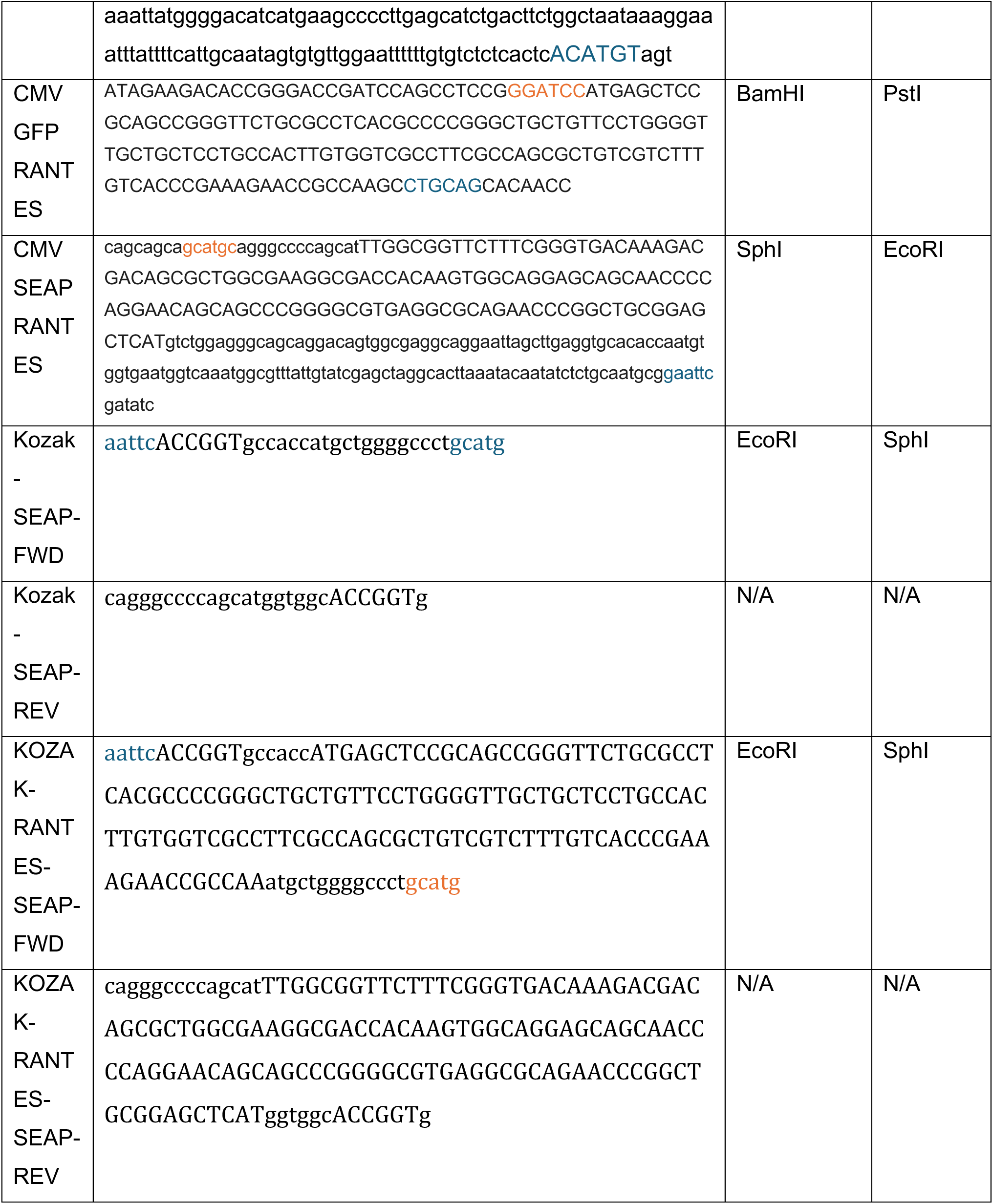

